## Supplementary figures and images for "Phenotypic and molecular evolution across 10,000 generations in laboratory budding yeast populations"

### cropped_P1B02.png

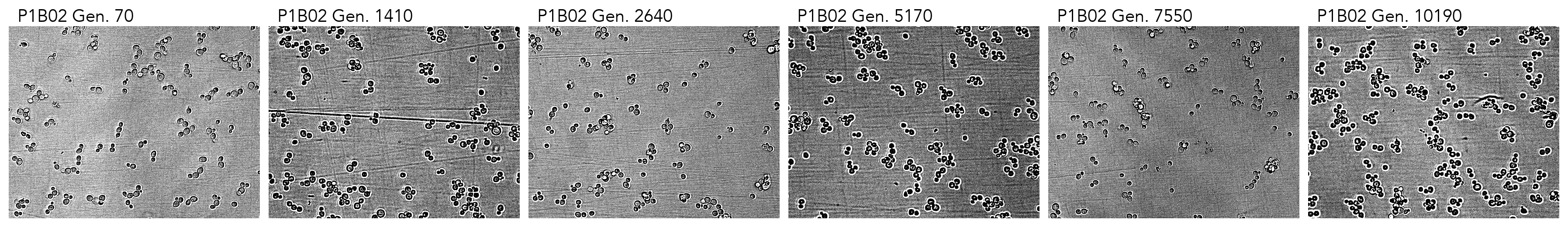

### cropped_P1B03.png

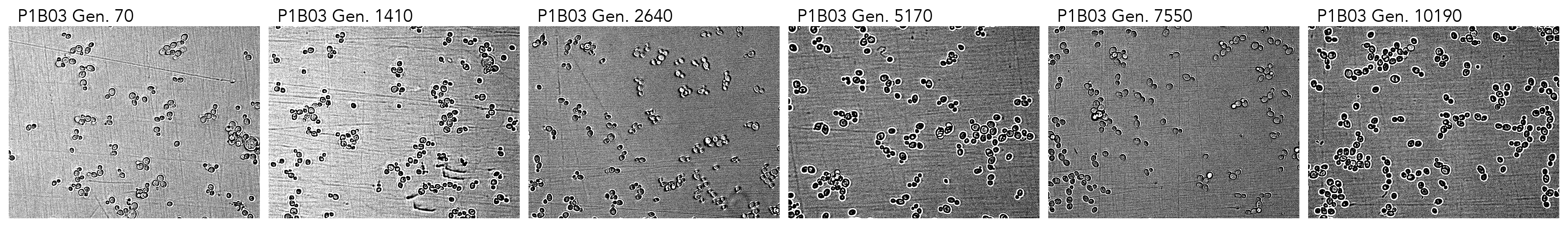

### cropped_P1B04.png

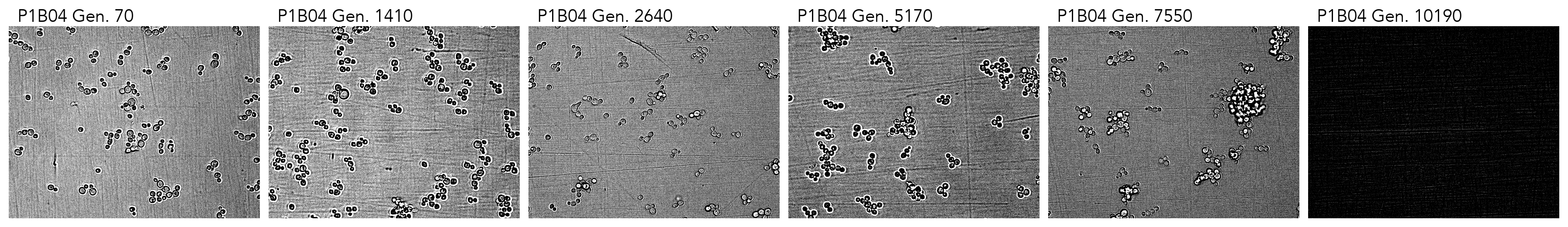

### cropped_P1B07.png

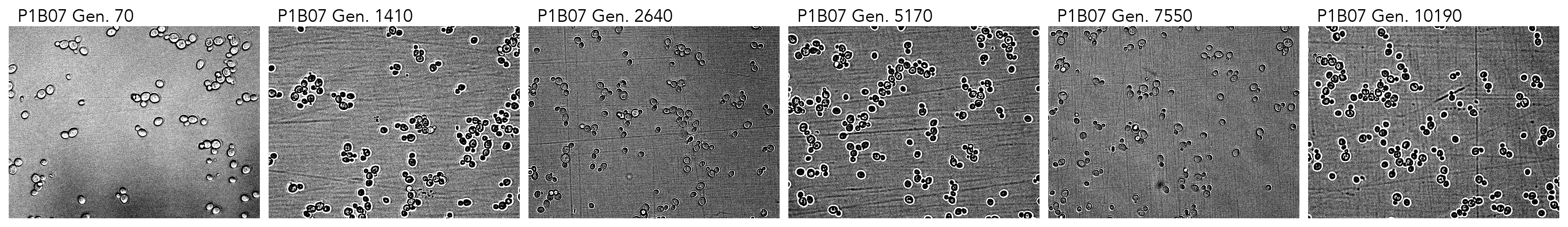

### cropped_P1B11.png

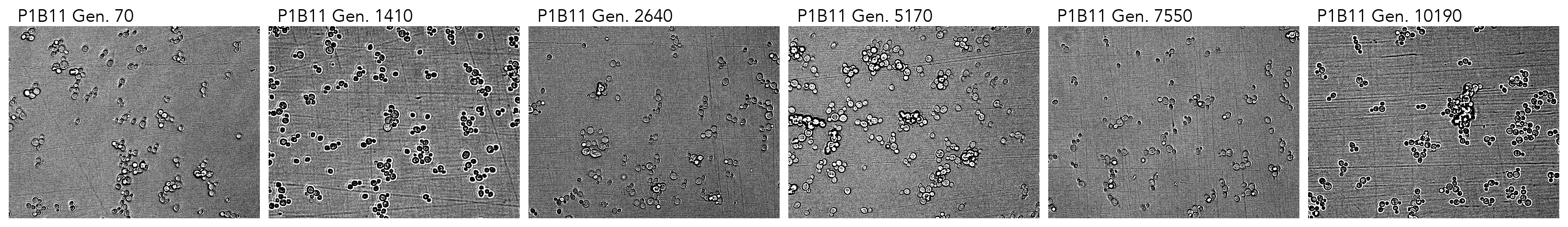

### cropped_P1C02.png

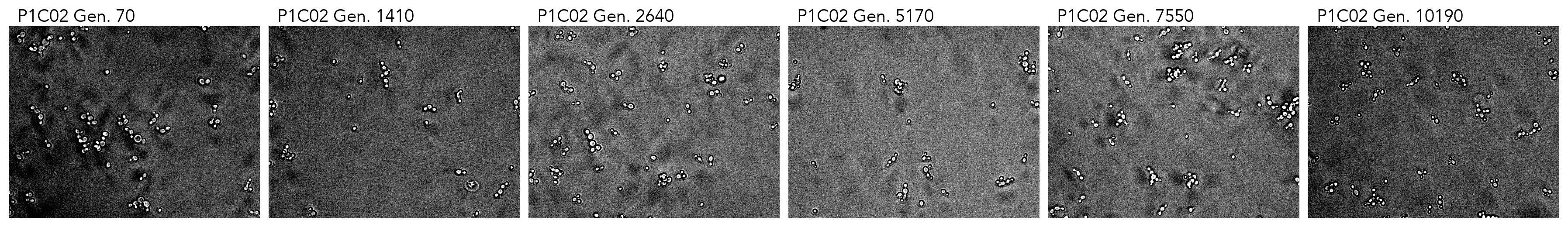

### cropped_P1C04.png

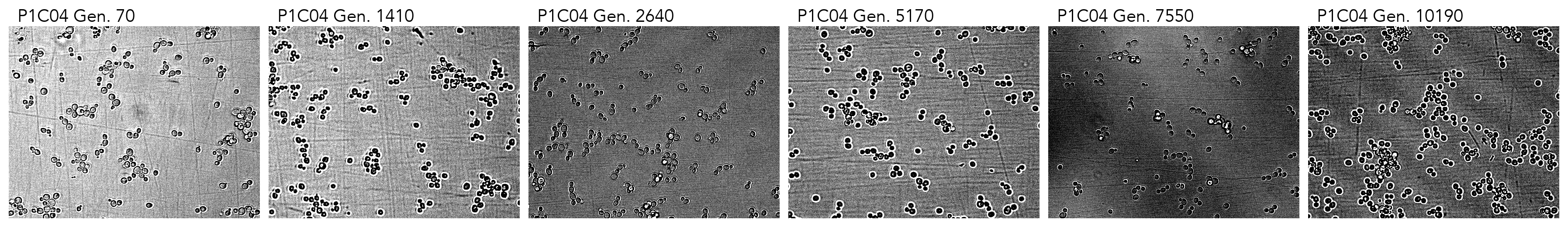

### cropped_P1C05.png

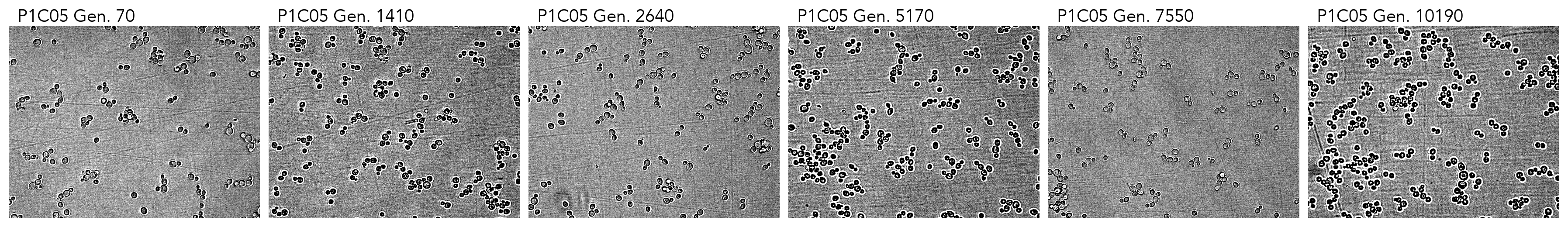

### cropped_P1C06.png

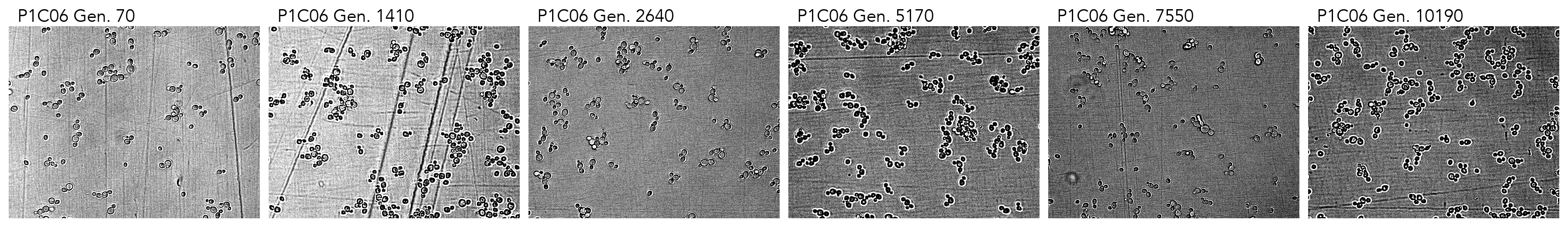

### cropped_P1C07.png

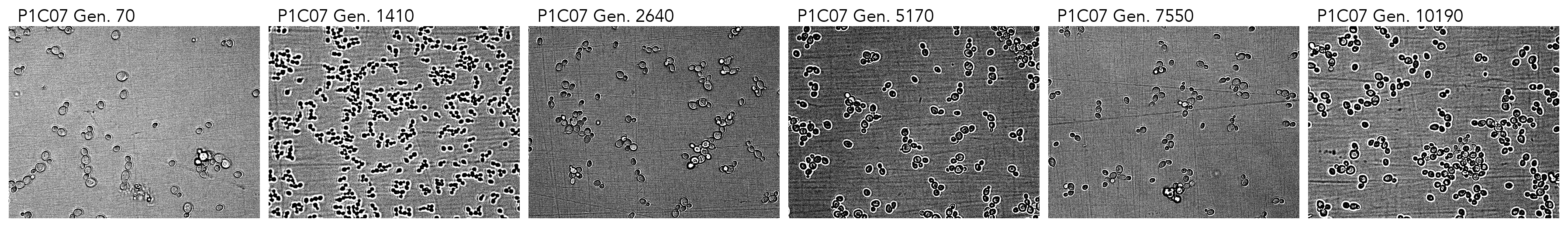

### cropped_P1C08.png

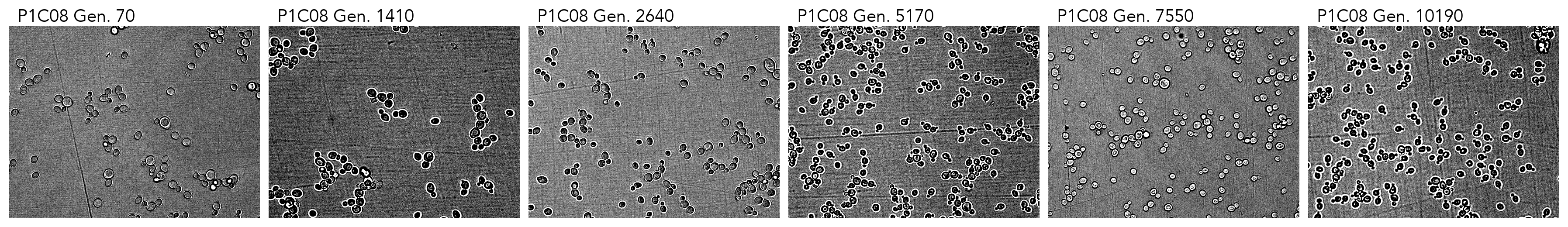

### cropped_P1C09.png

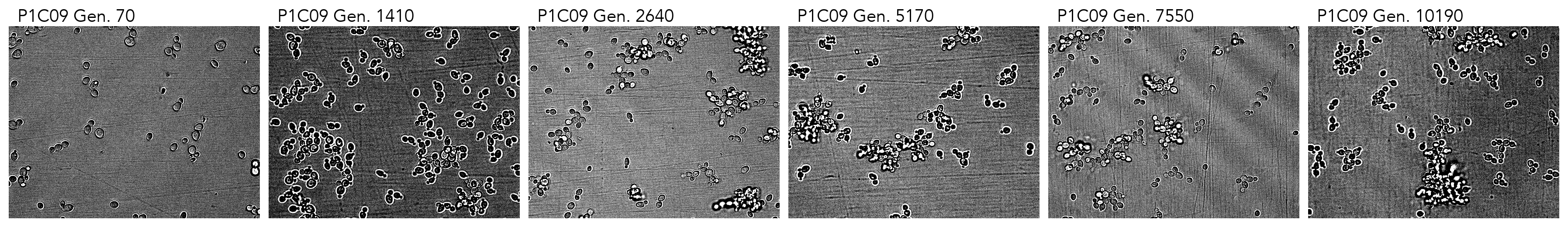

### cropped_P1C11.png

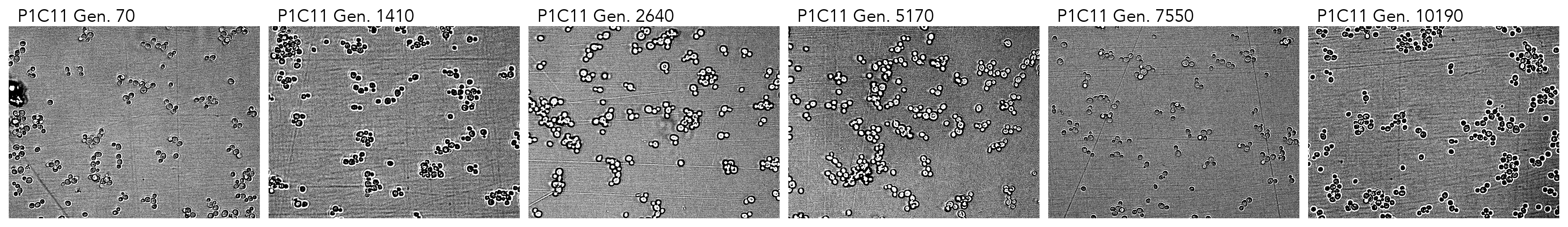

### cropped_P1D03.png

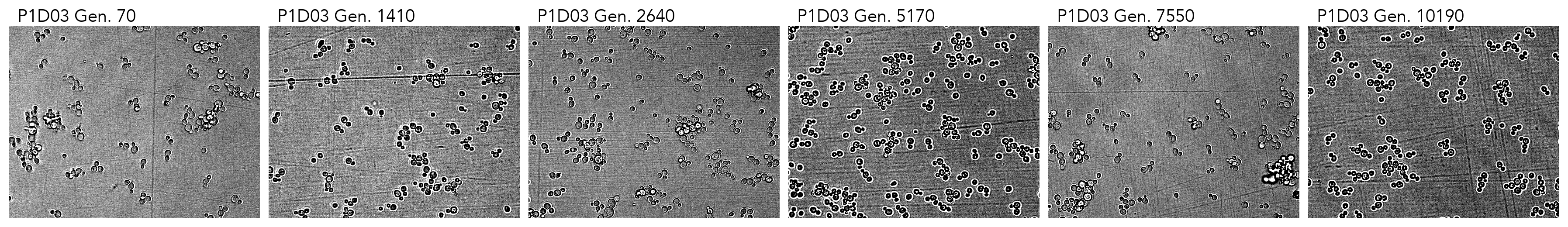

### cropped_P1D09.png

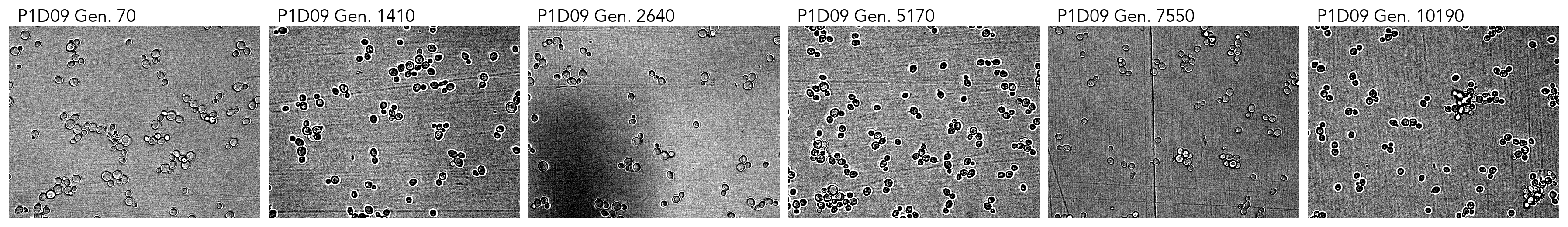

### cropped_P1E04.png

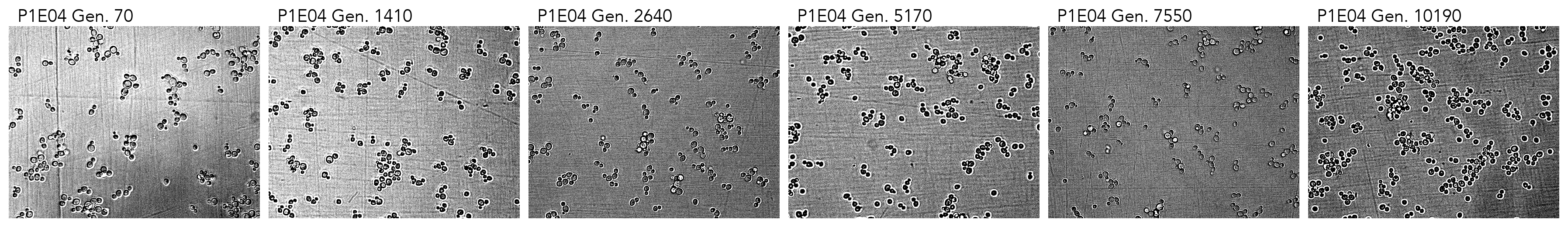

### cropped_P1E09.png

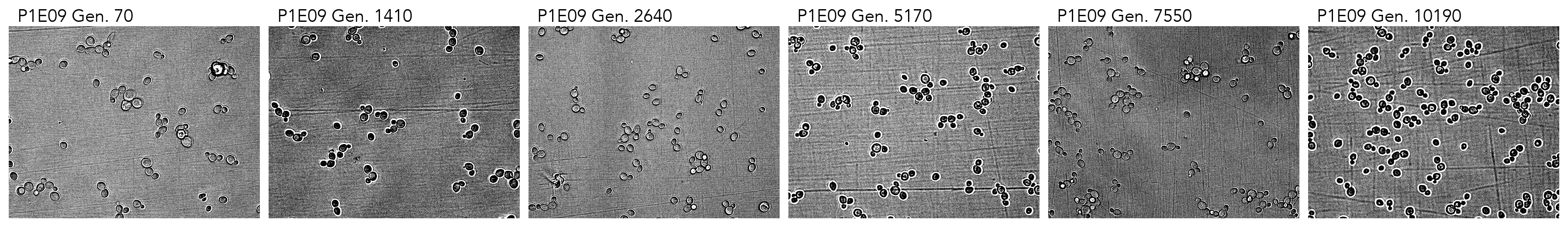

### cropped_P1E11.png

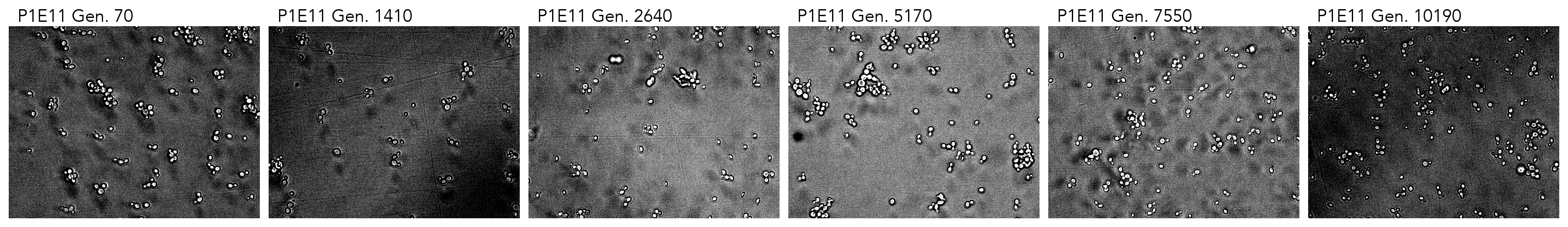

### cropped_P1F05.png

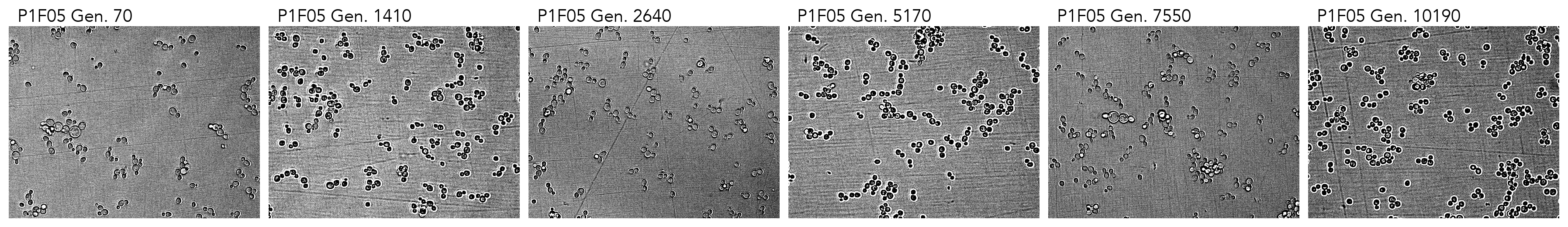

### cropped_P1F07.png

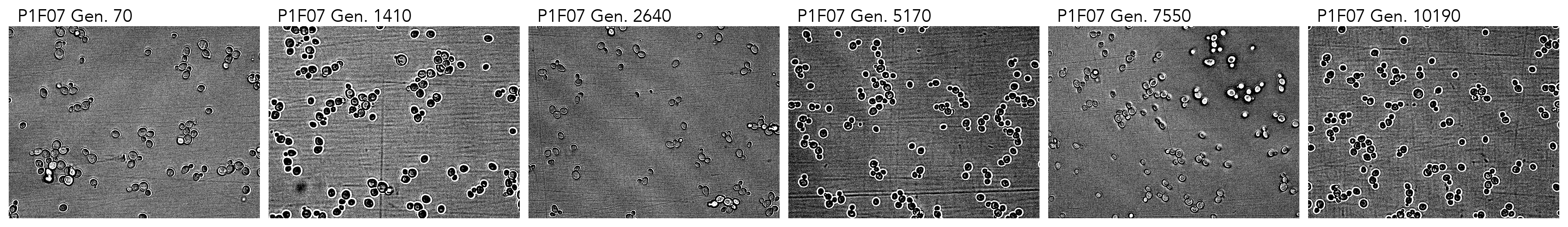

### cropped_P1F08.png

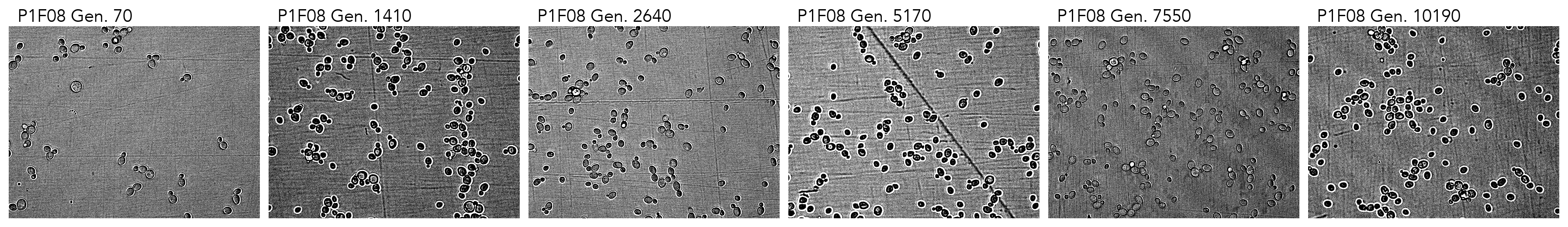

### cropped_P1F10.png

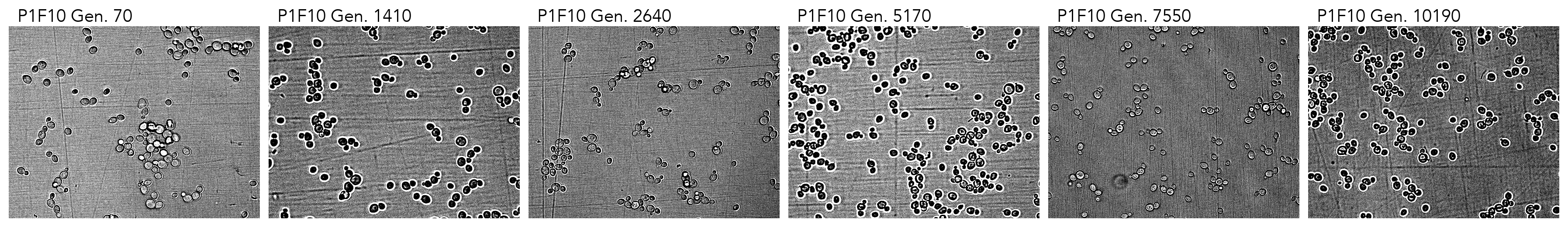

### cropped_P1F11.png

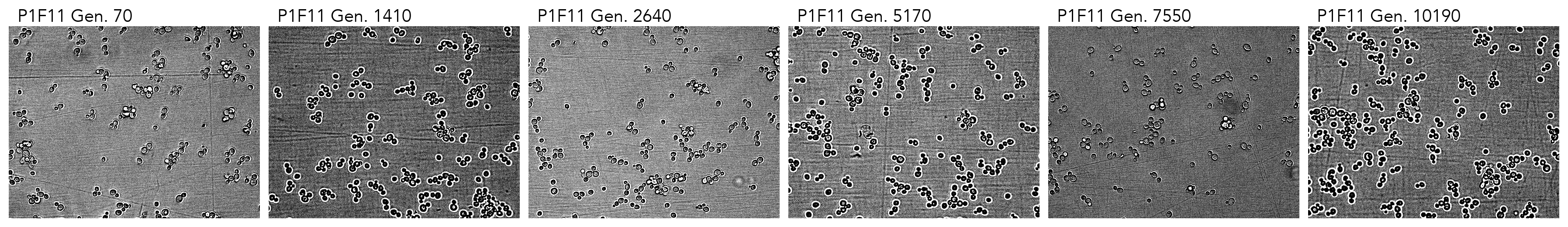

### cropped_P1G04.png

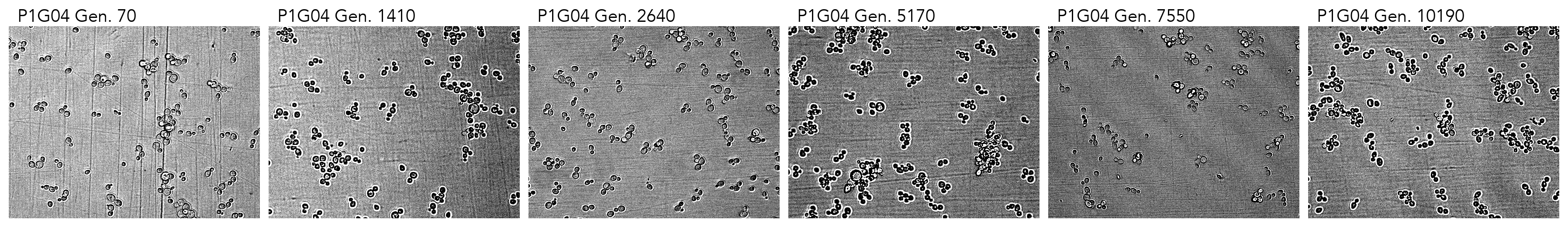

### cropped_P1G05.png

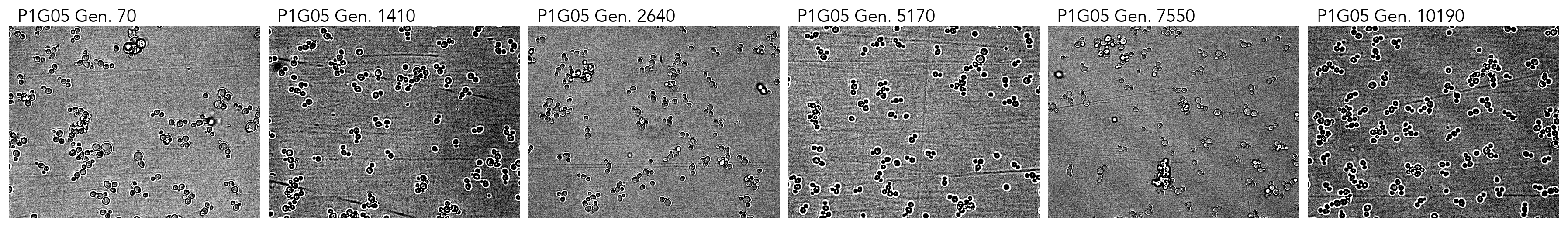

### cropped_P1G08.png

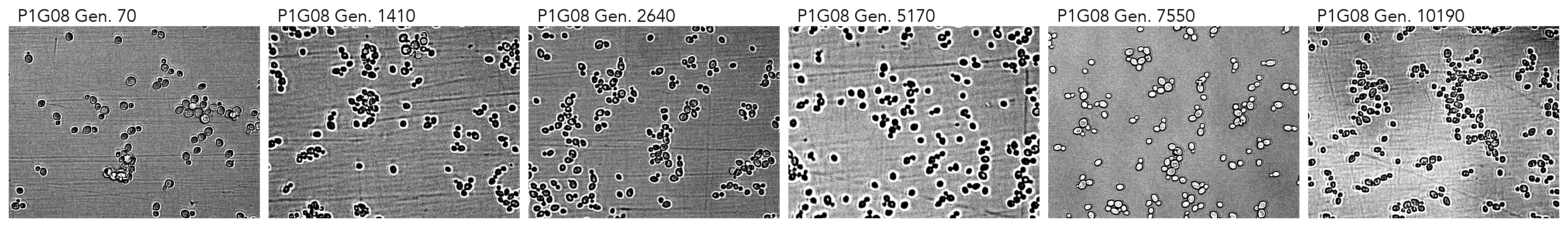

### cropped_P1G09.png

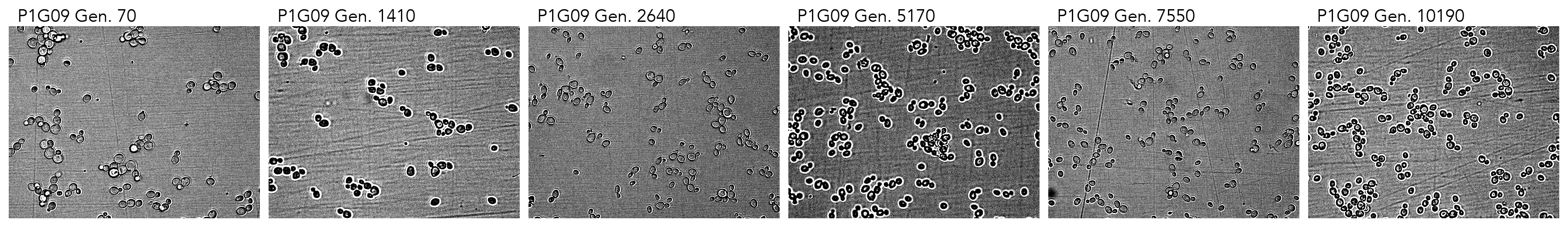

### cropped_P1G10.png

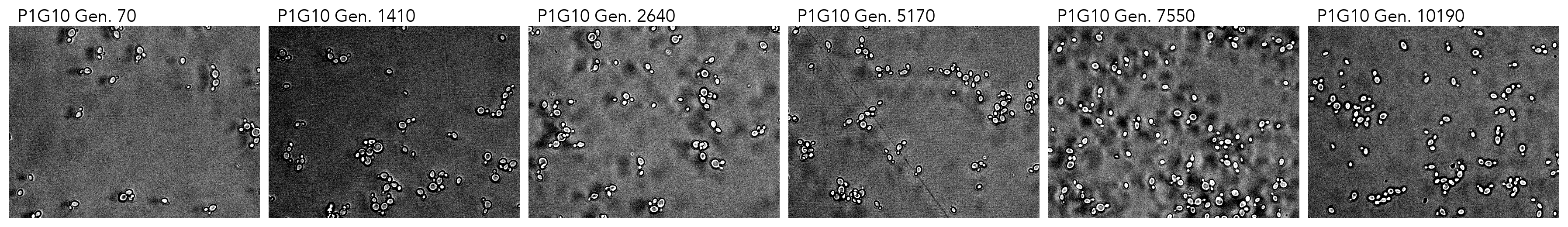

### cropped_P1G11.png

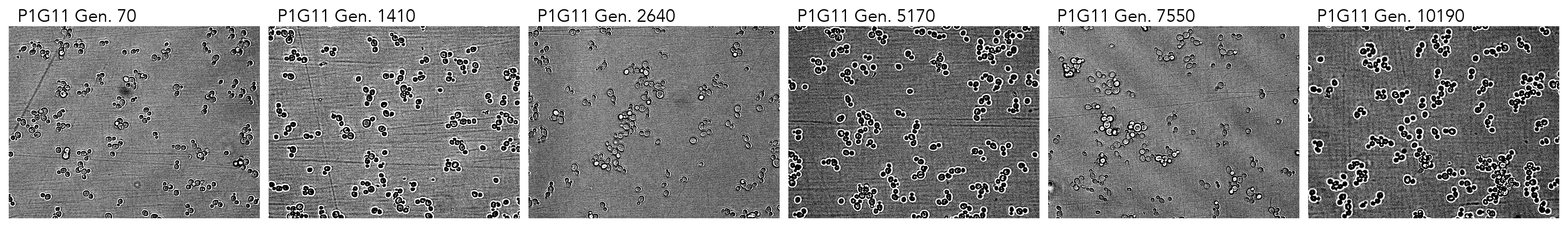

### cropped_P1H11.png

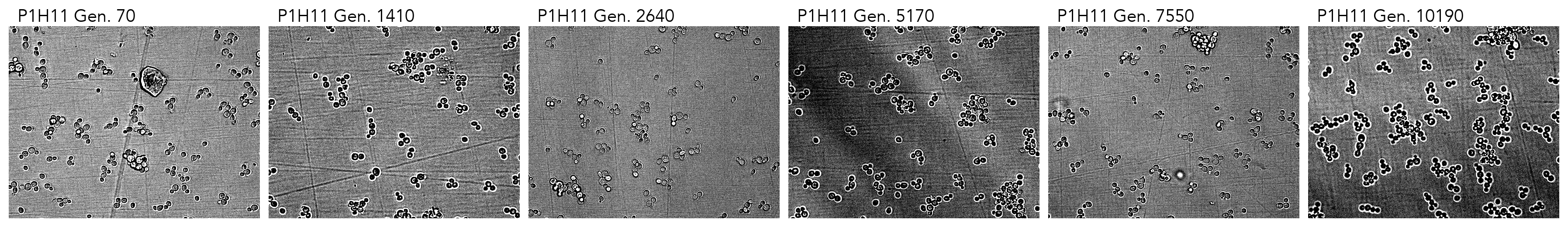

### cropped_P2B04.png

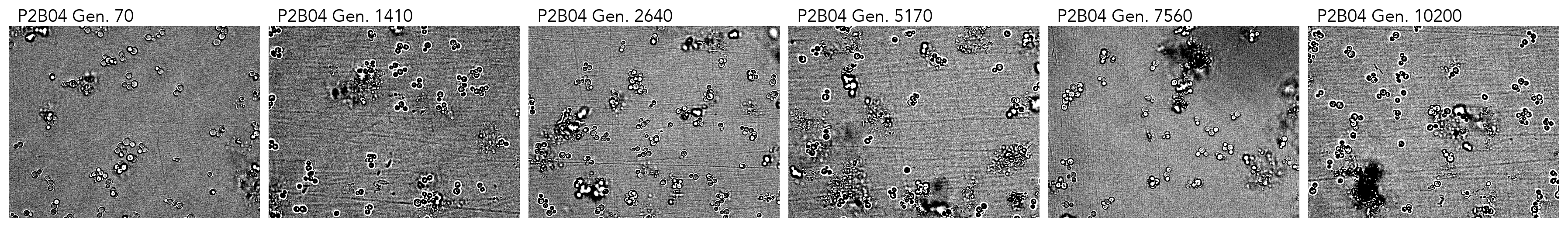

### cropped_P2B05.png

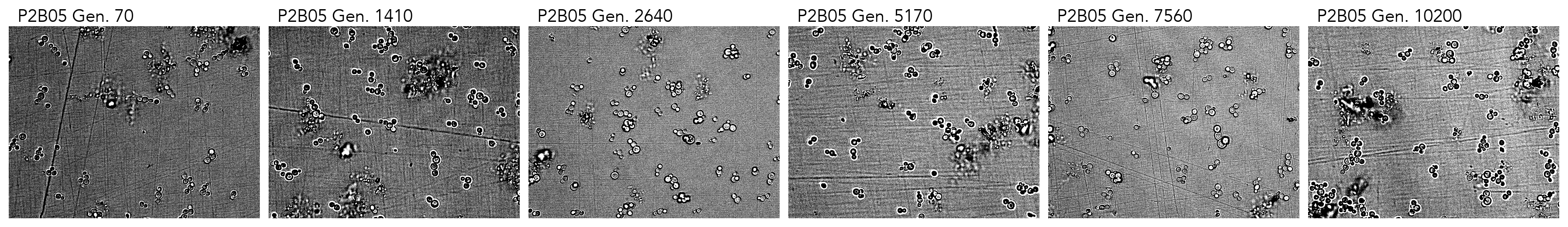

### cropped_P2B07.png

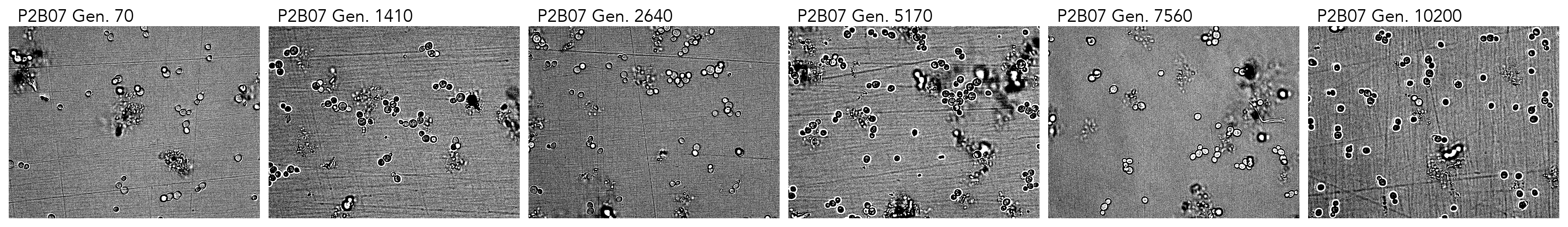

### cropped_P2B08.png

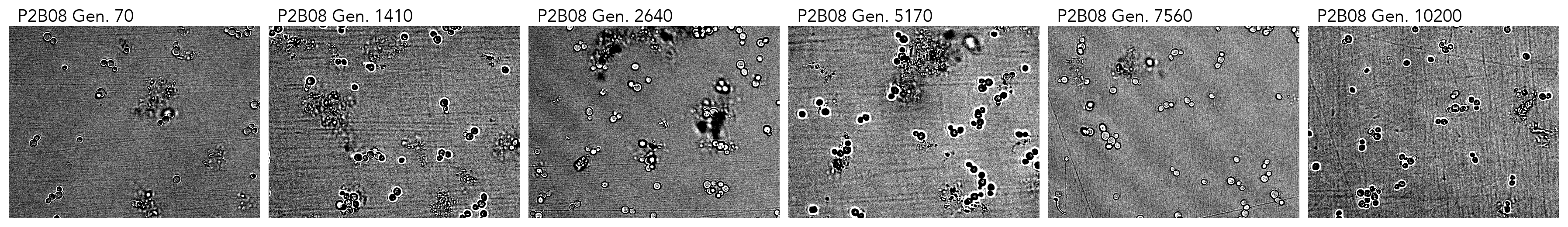

### cropped_P2B09.png

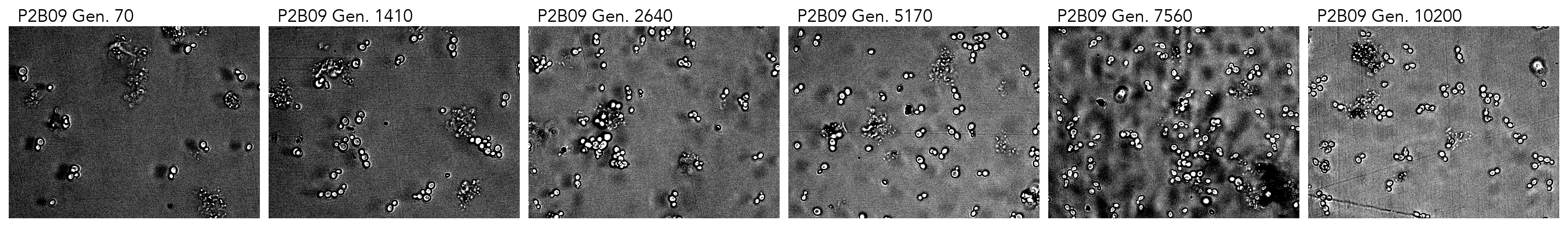

### cropped_P2B10.png

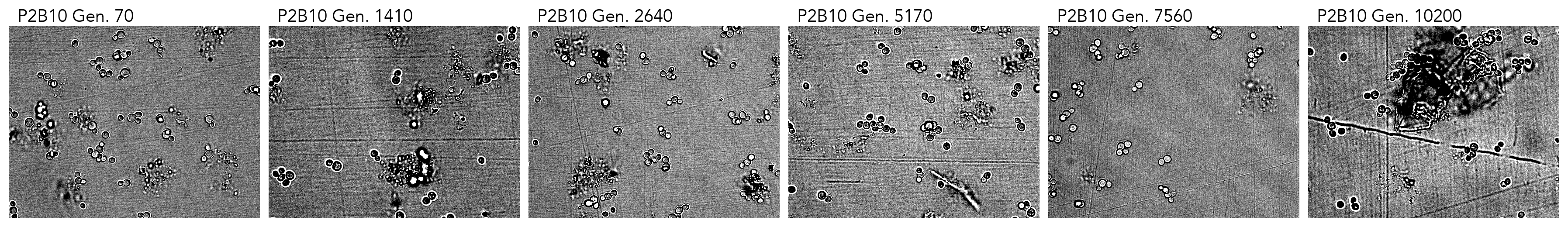

### cropped_P2B11.png

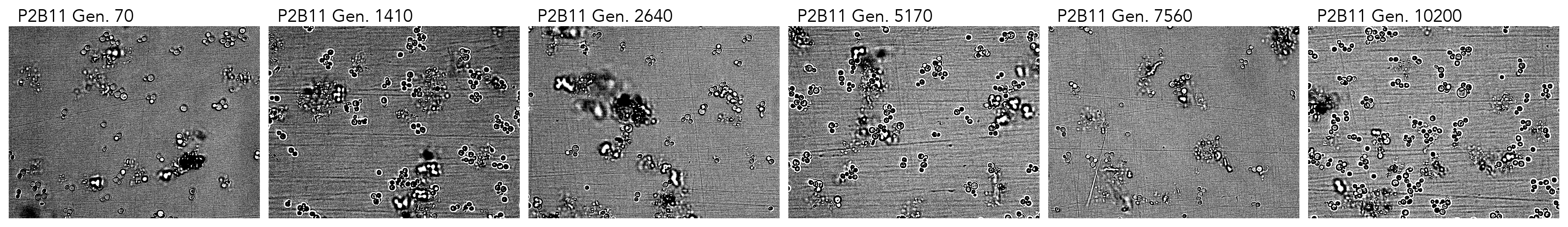

### cropped_P2C02.png

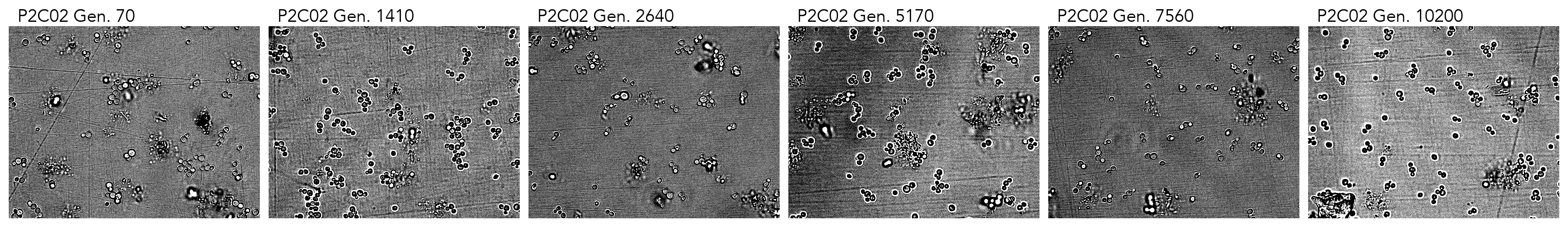

### cropped_P2C04.png

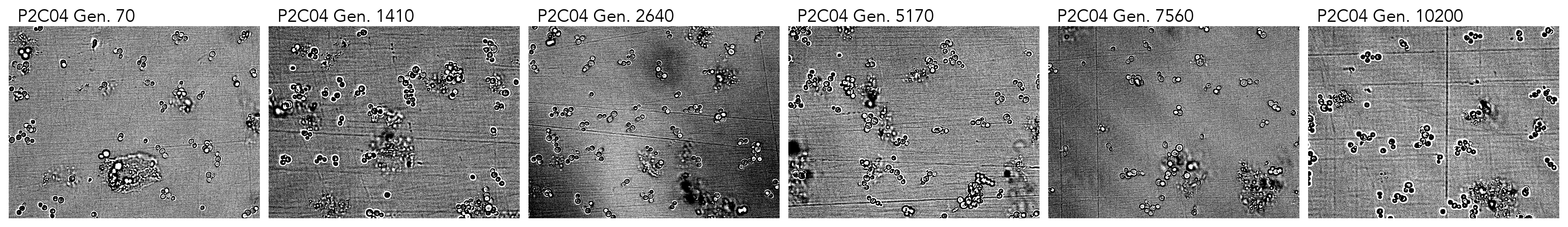

### cropped_P2C05.png

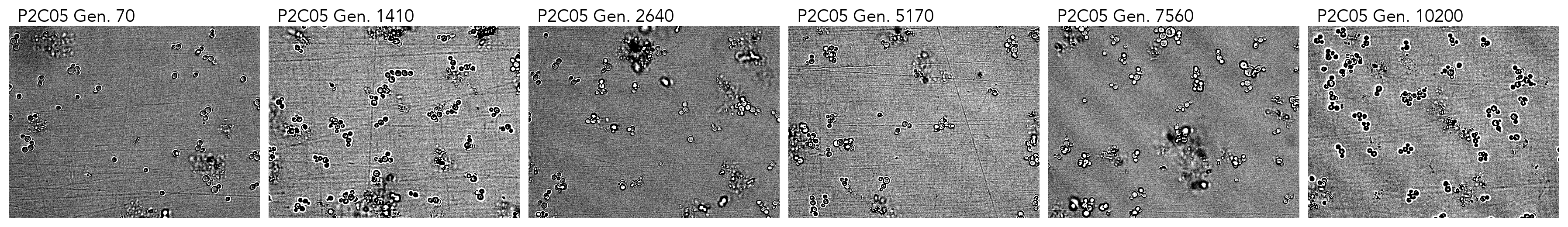

### cropped_P2C06.png

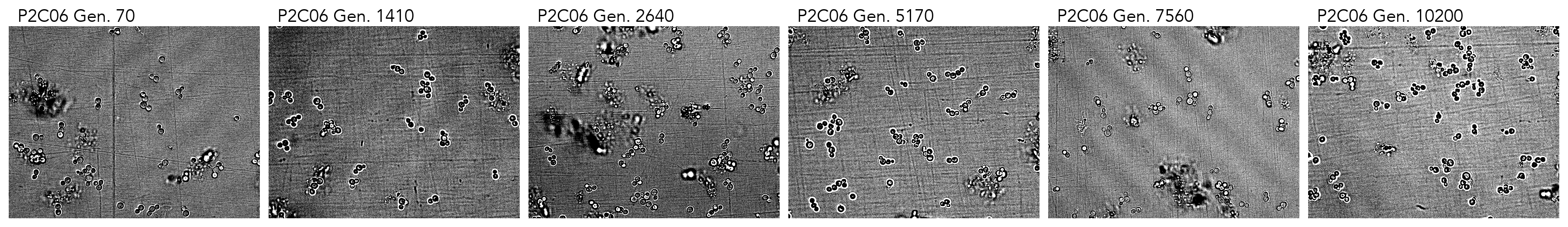

### cropped_P2C10.png

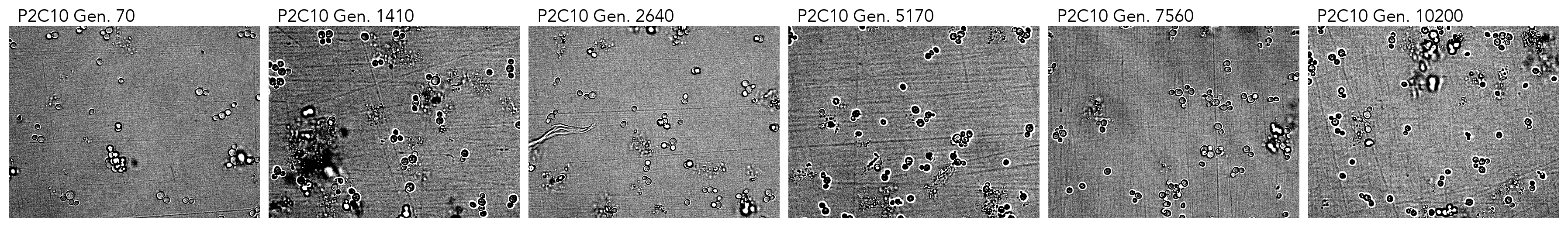

### cropped_P2C11.png

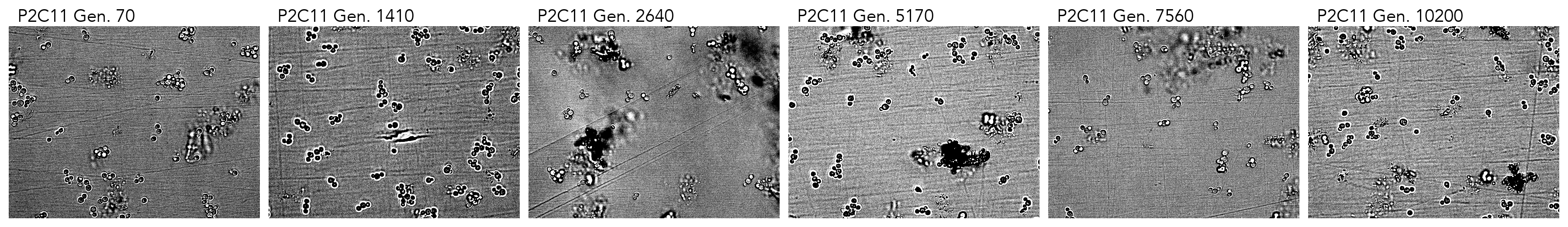

### cropped_P2D03.png

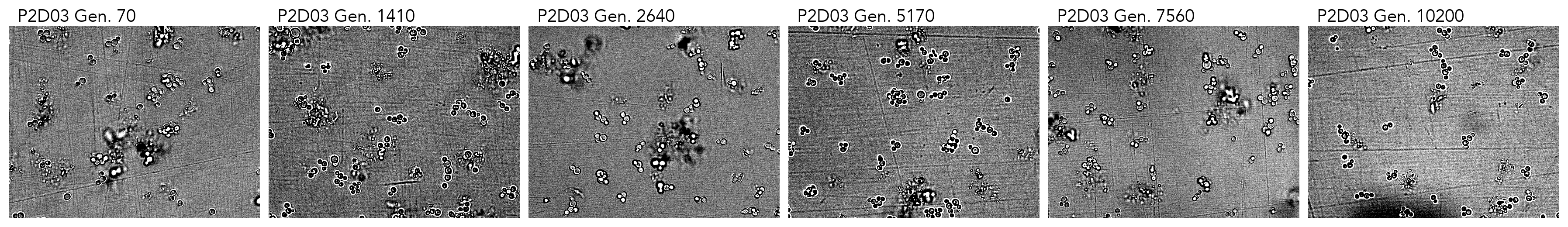

### cropped_P2D06.png

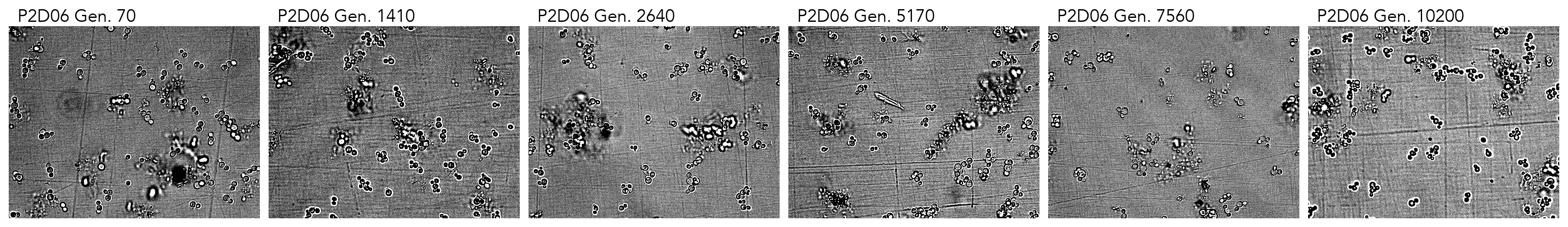

### cropped_P2D08.png

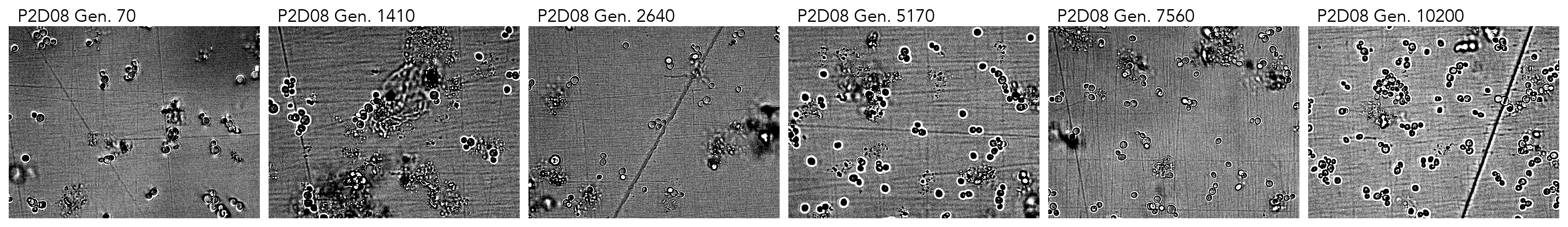

### cropped_P2D11.png

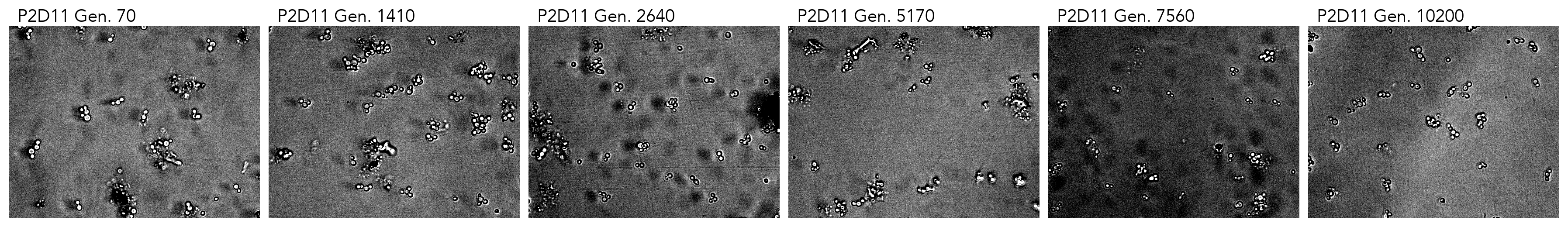

### cropped_P2E06.png

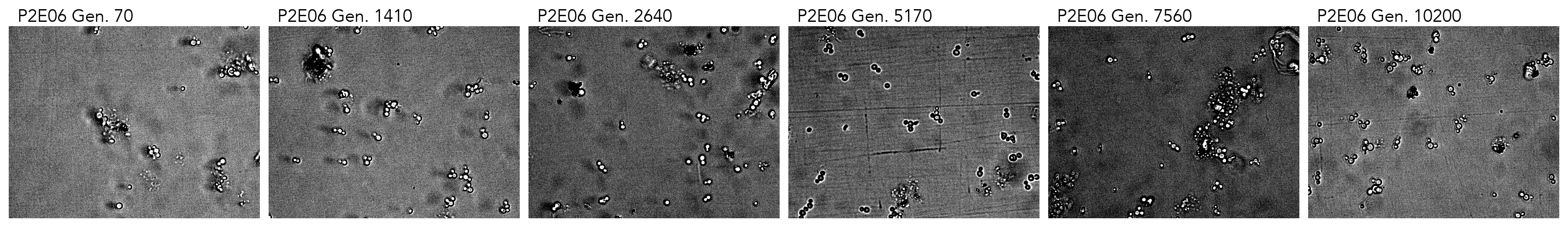

### cropped_P2E08.png

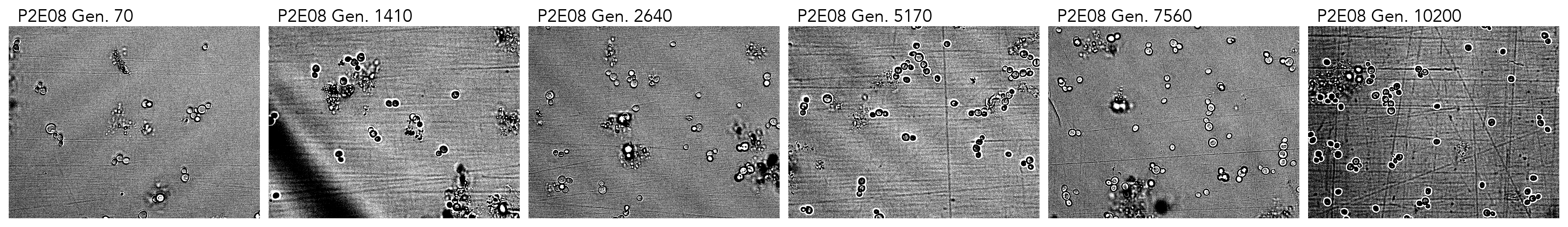

### cropped_P2E11.png

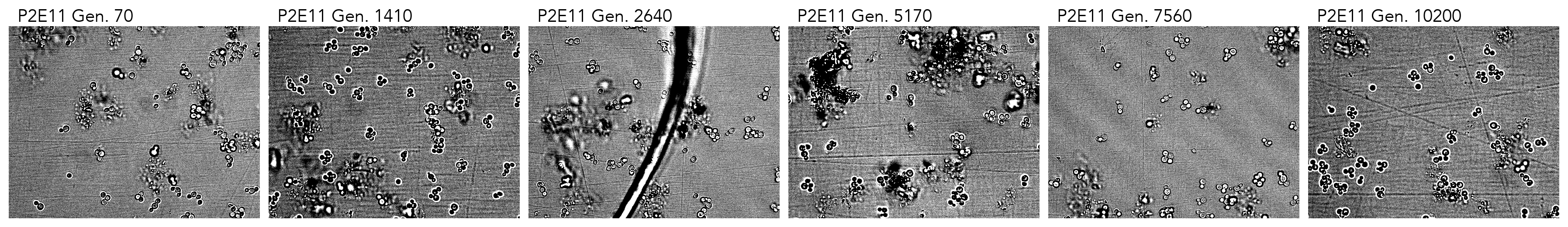

### cropped_P2F02.png

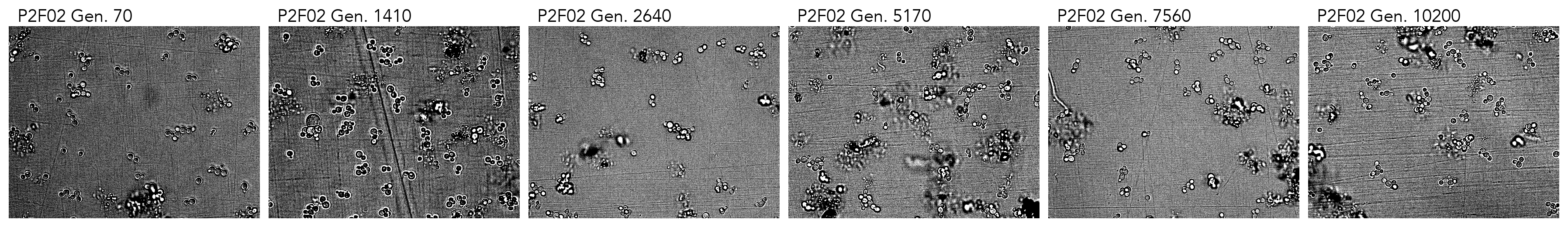

### cropped_P2F07.png

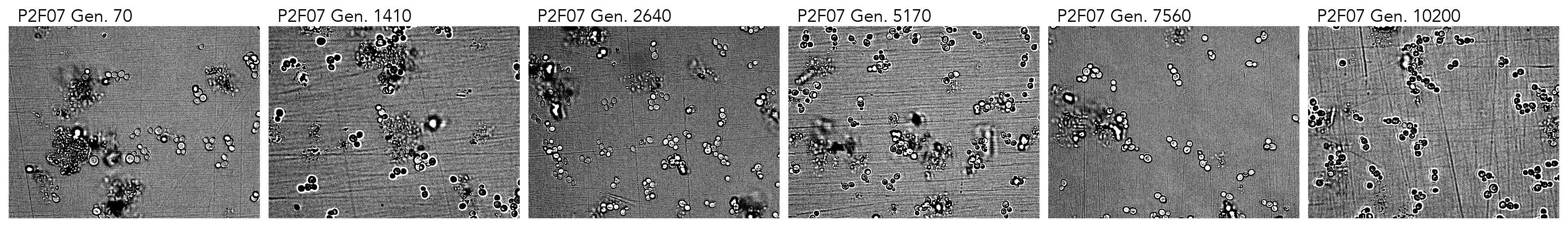

### cropped_P2F09.png

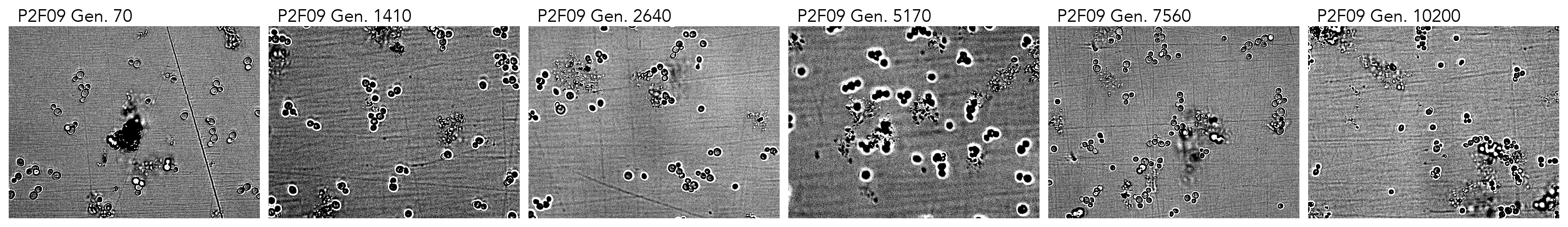

### cropped_P2F11.png

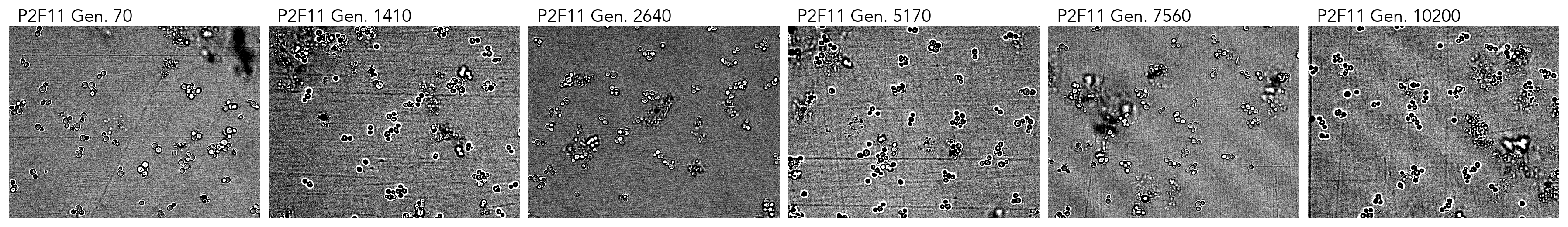

### cropped_P2G04.png

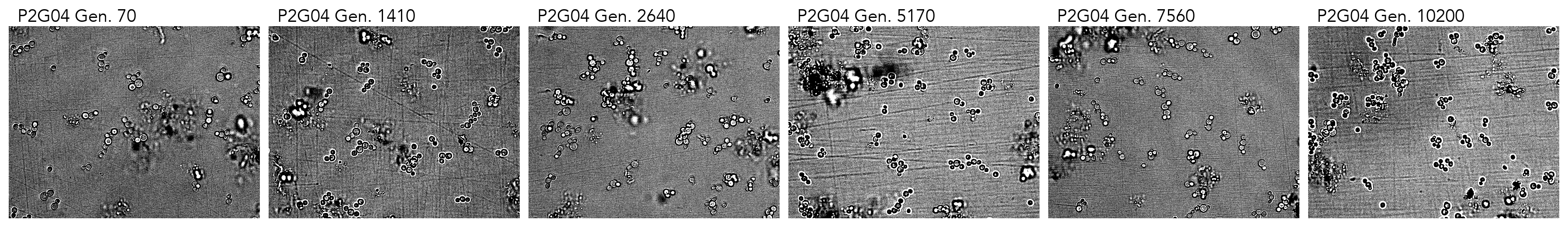

### cropped_P2G05.png

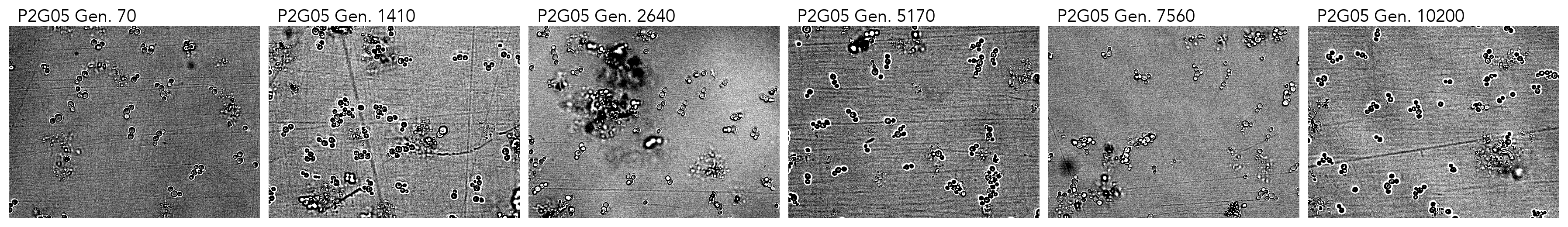

### cropped_P2G08.png

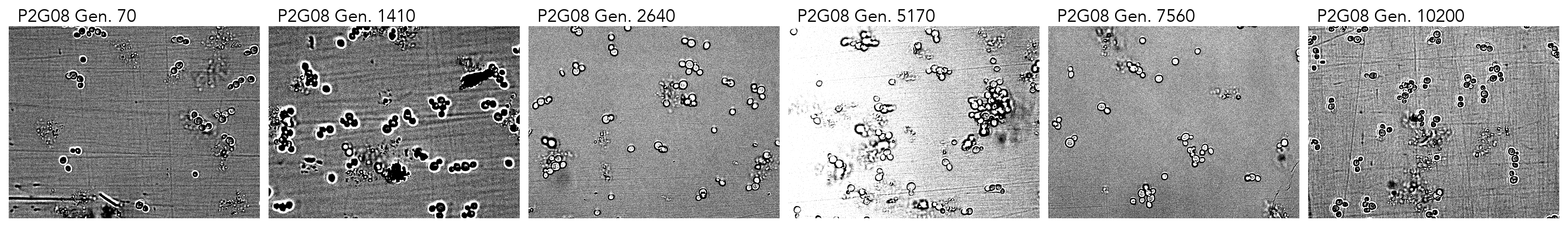

### cropped_P2G09.png

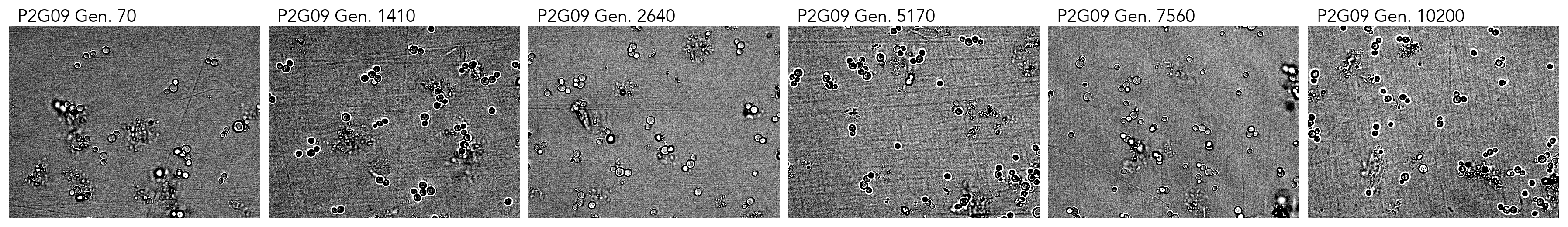

### cropped_P2G10.png

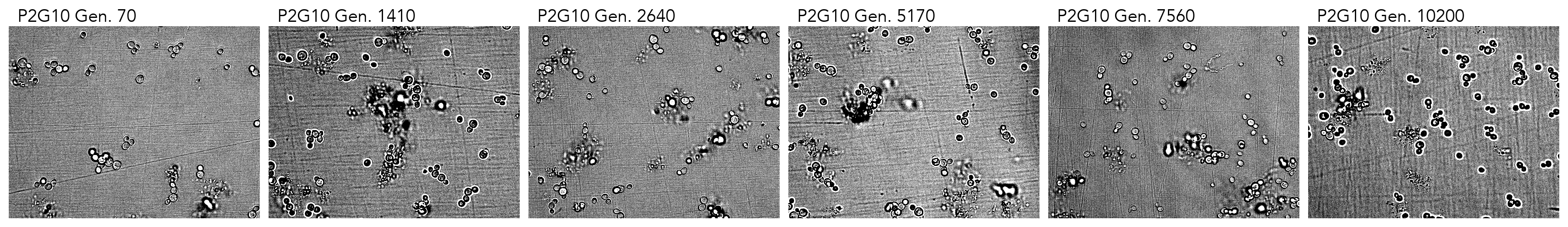

### cropped_P2G11.png

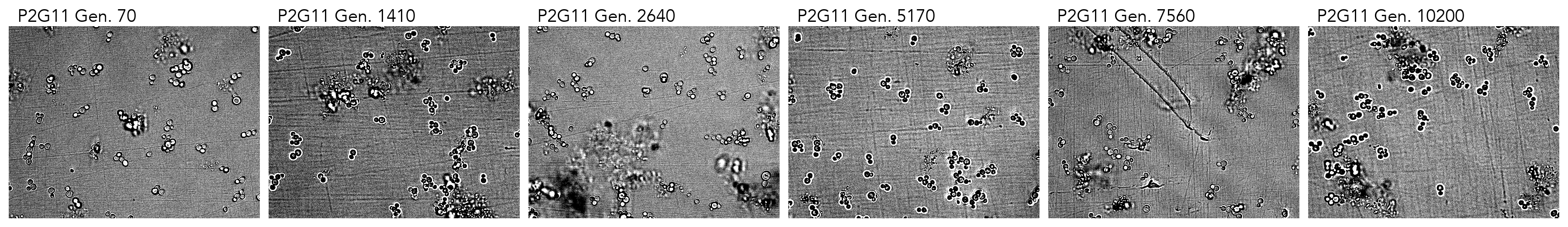

### cropped_P3B07.png

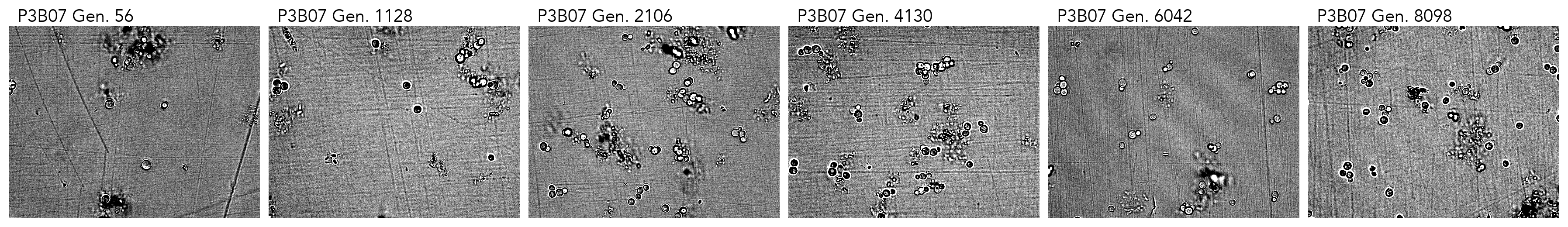

### cropped_P3B08.png

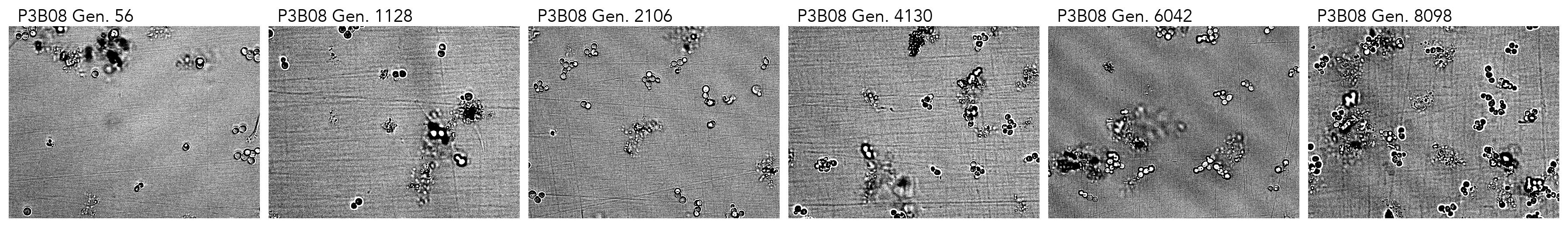

### cropped_P3B10.png

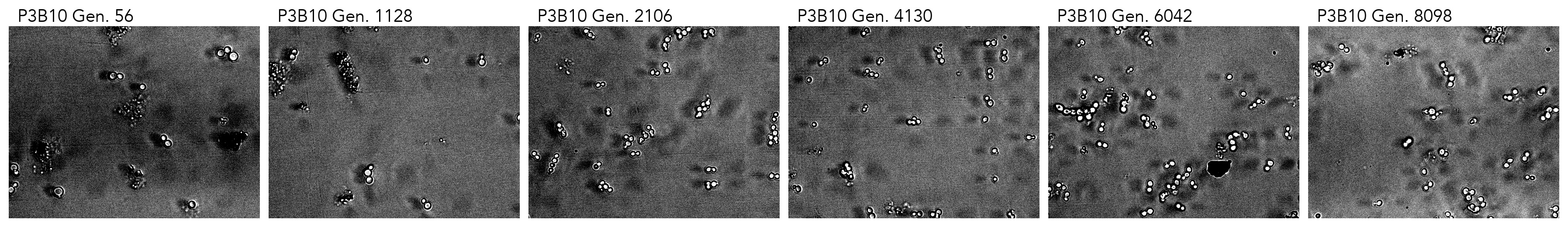

### cropped_P3B11.png

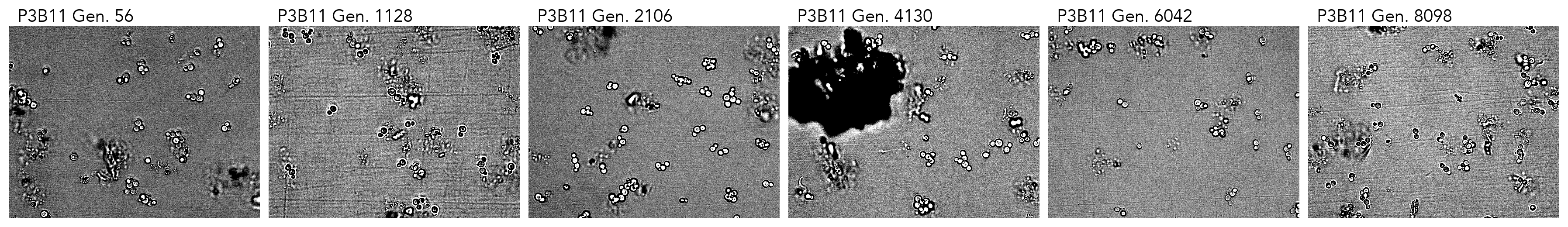

### cropped_P3C03.png

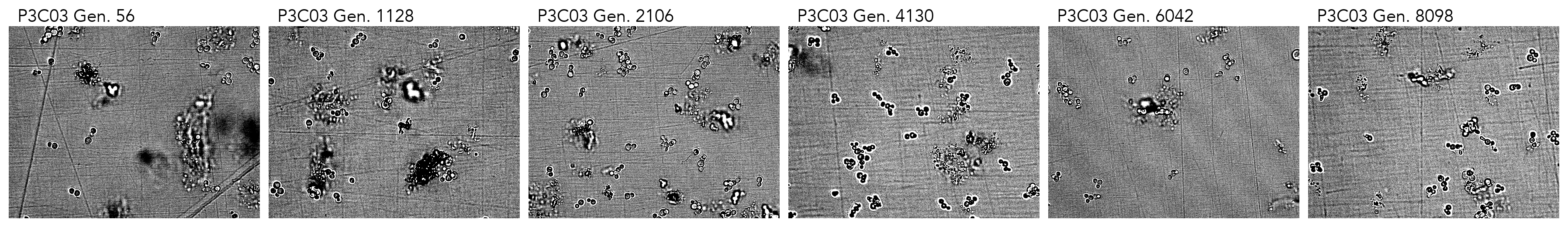

### cropped_P3C04.png

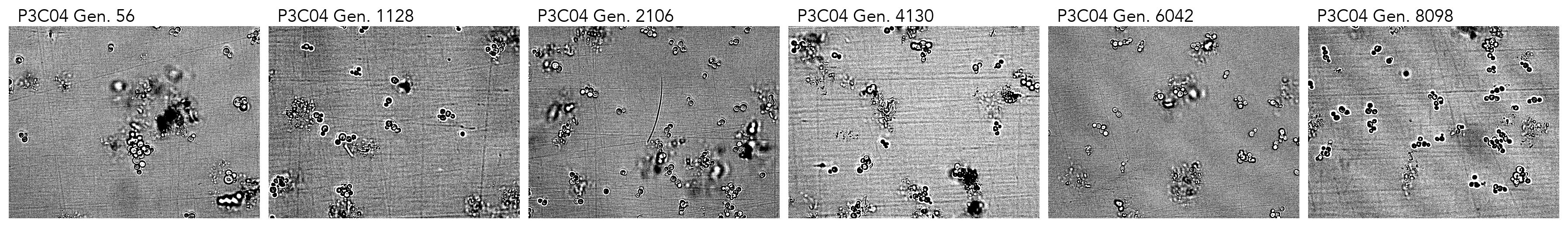

### cropped_P3C05.png

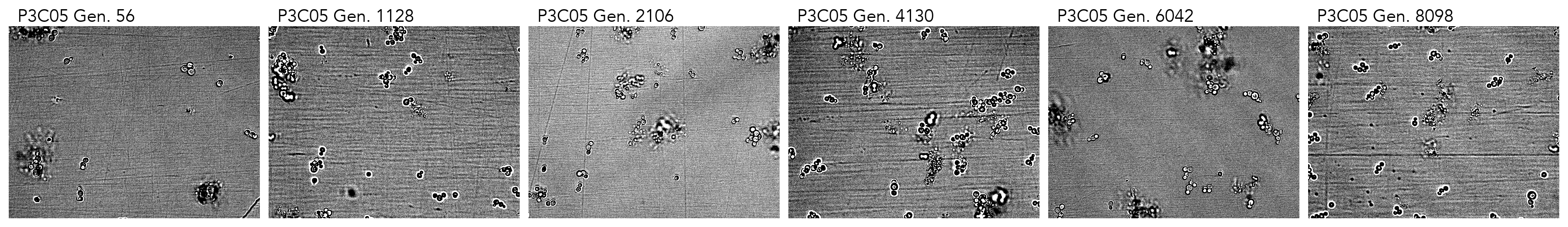

### cropped_P3C07.png

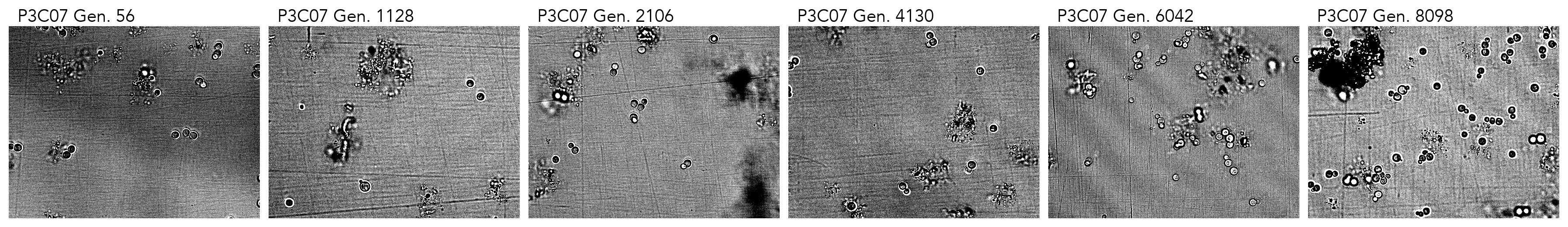

### cropped_P3C10.png

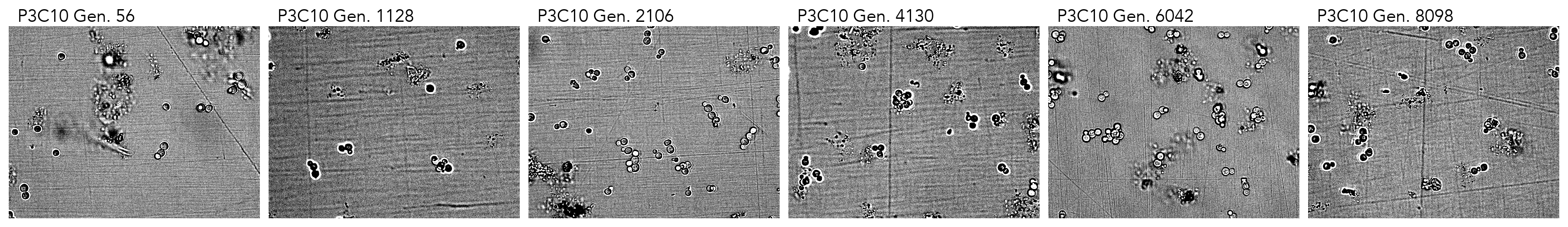

### cropped_P3C11.png

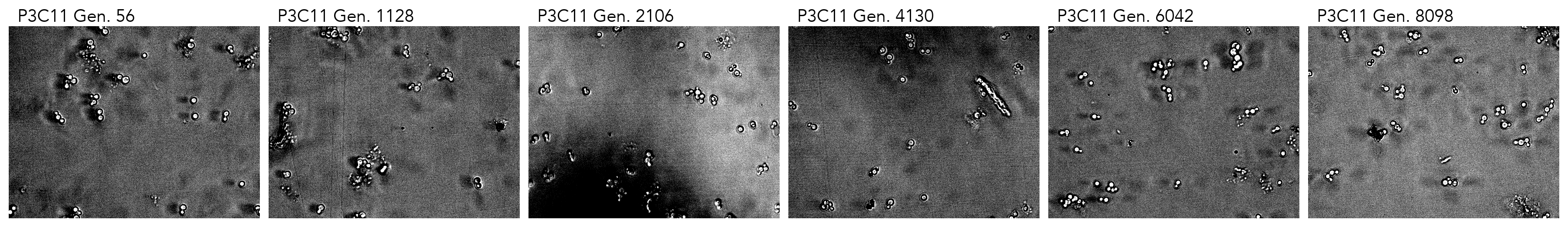

### cropped_P3D02.png

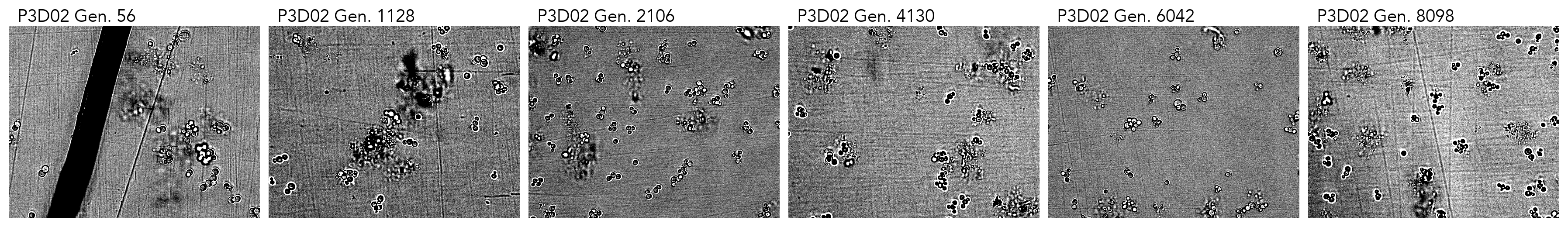

### cropped_P3D03.png

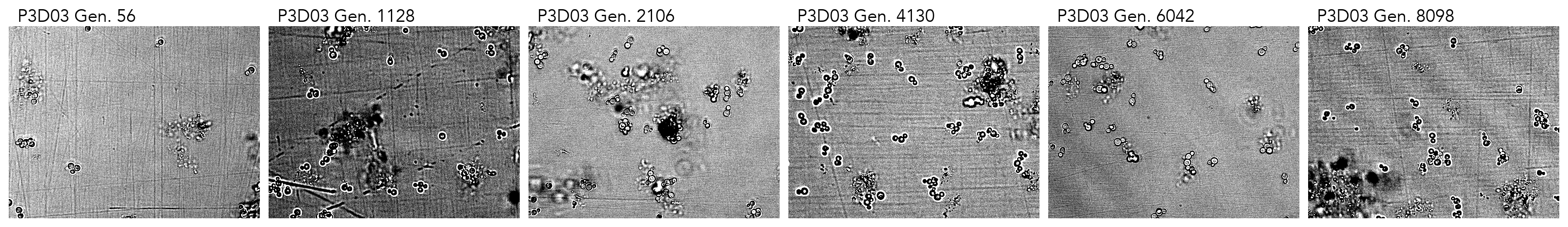

### cropped_P3D05.png

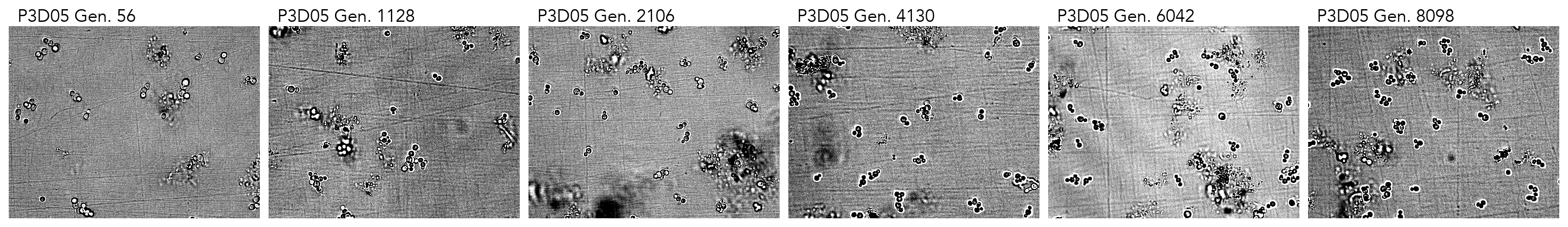

### cropped_P3D09.png

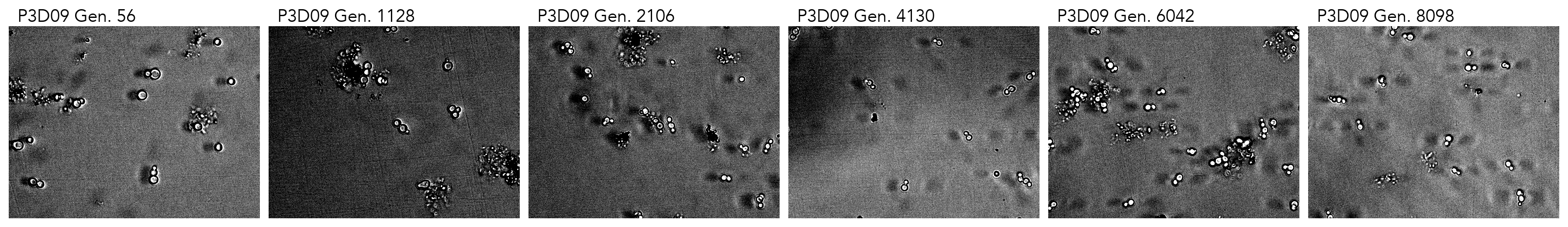

### cropped_P3D10.png

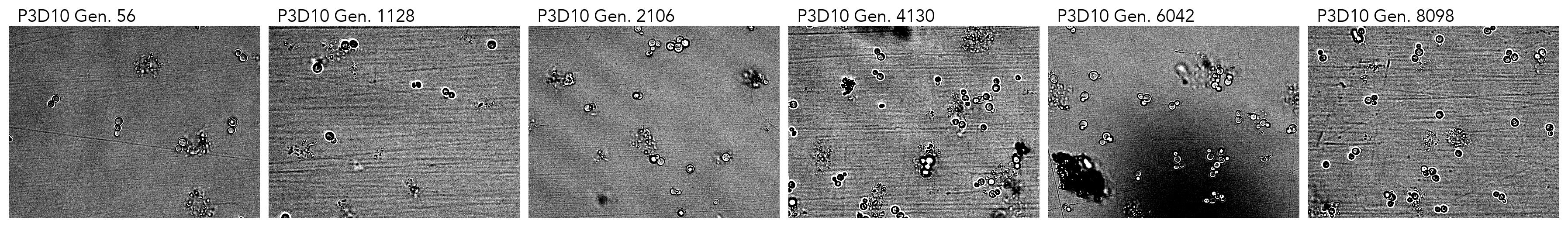

### cropped_P3D11.png

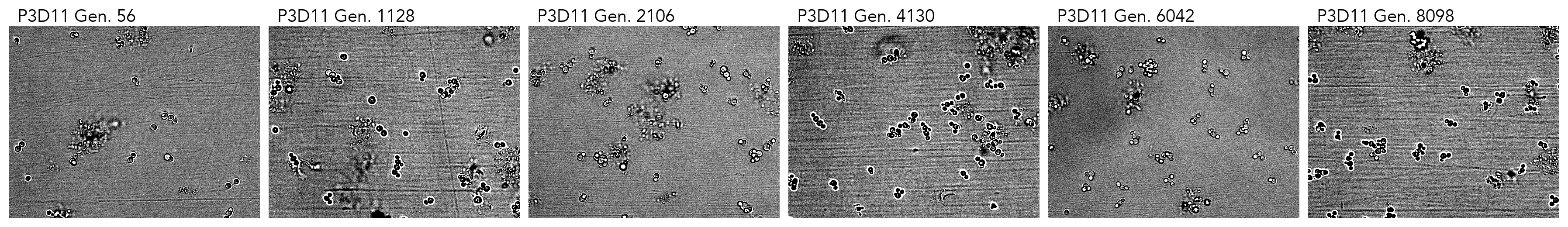

### cropped_P3E02.png

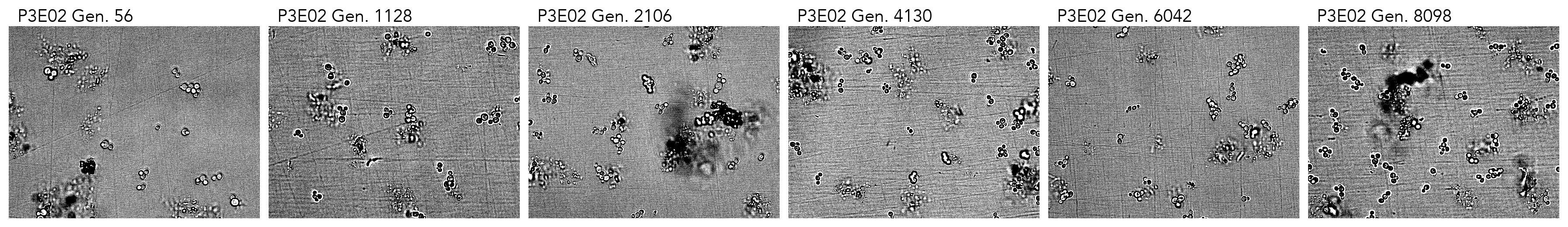

### cropped_P3E08.png

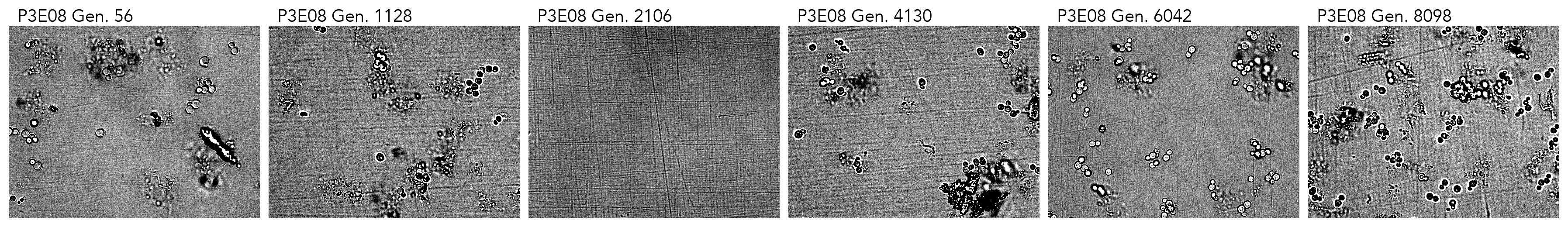

### cropped_P3E11.png

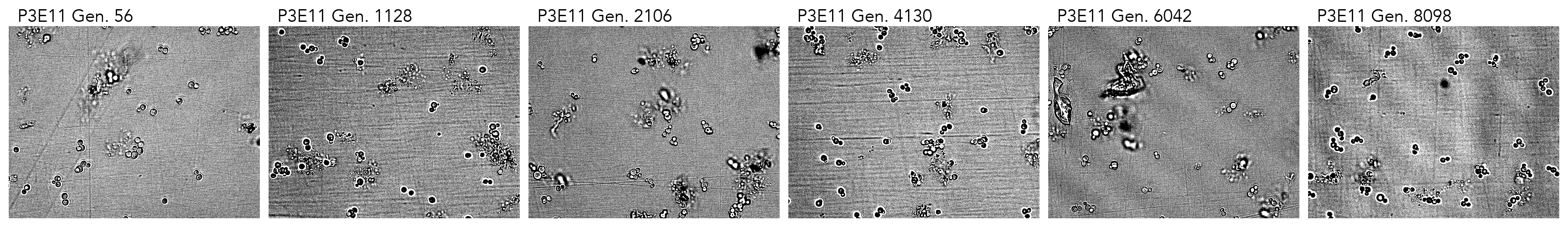

### cropped_P3F03.png

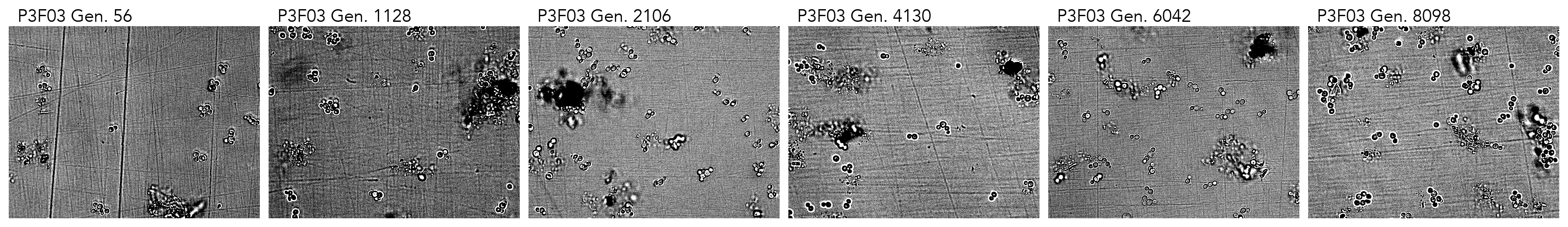

### cropped_P3F05.png

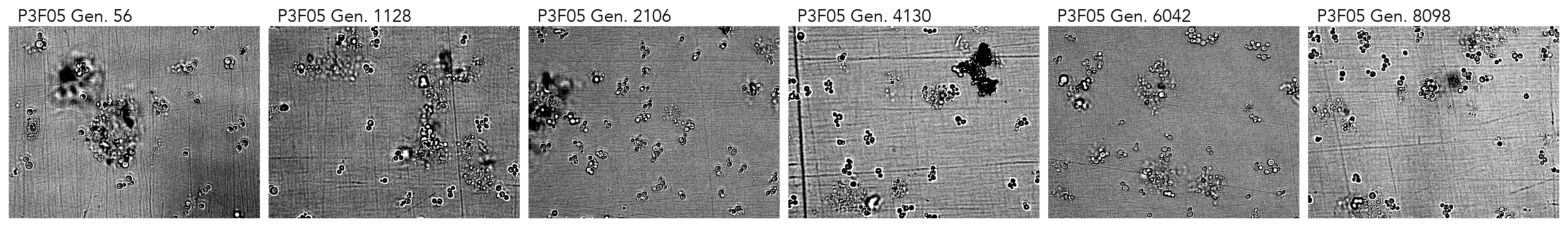

### cropped_P3F07.png

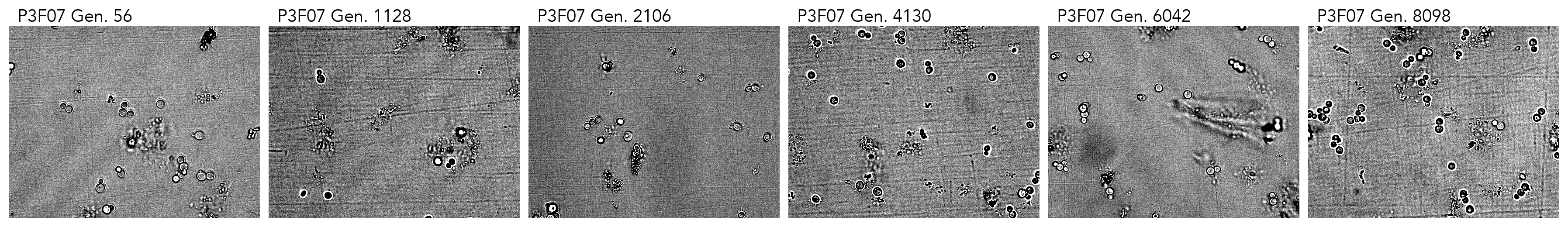

### cropped_P3F09.png

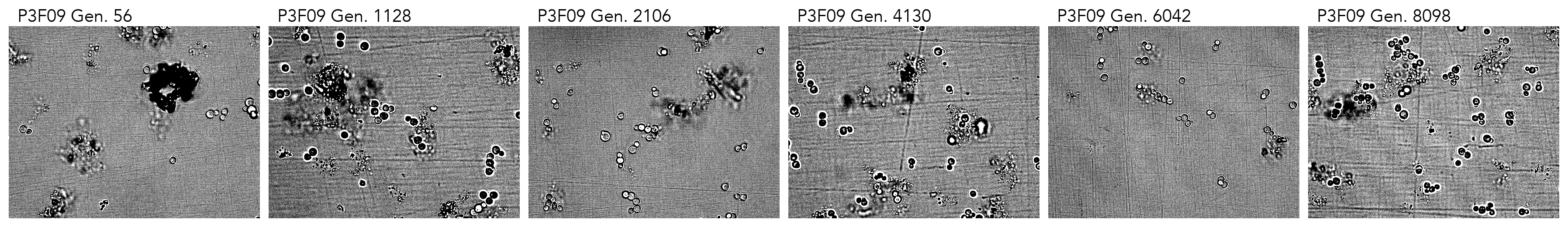

### cropped_P3F11.png

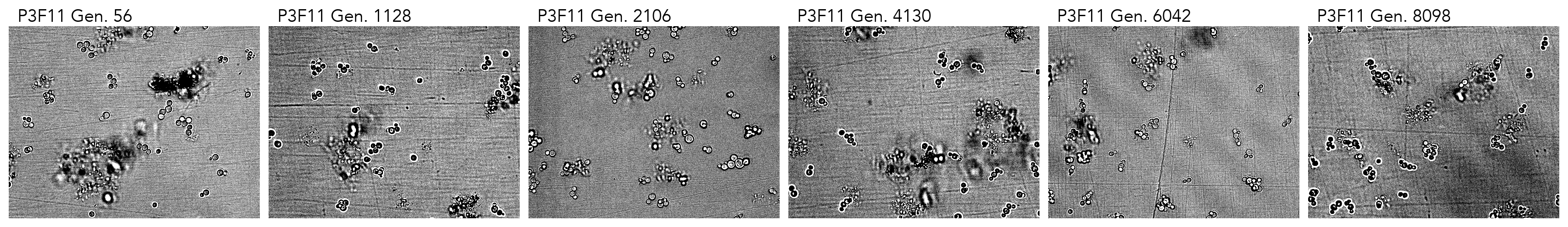

### cropped_P3G02.png

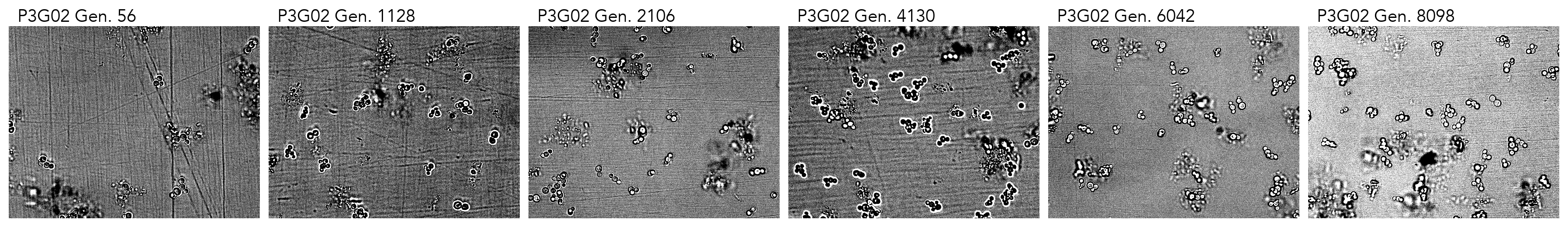

### cropped_P3G05.png

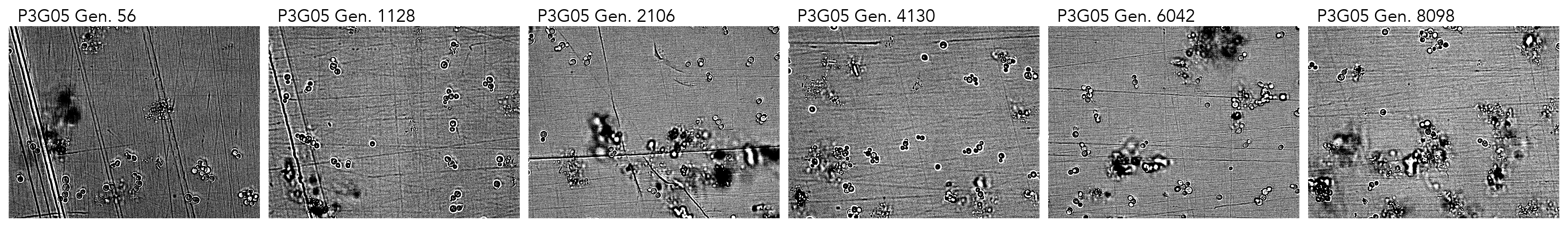

### cropped_P3G06.png

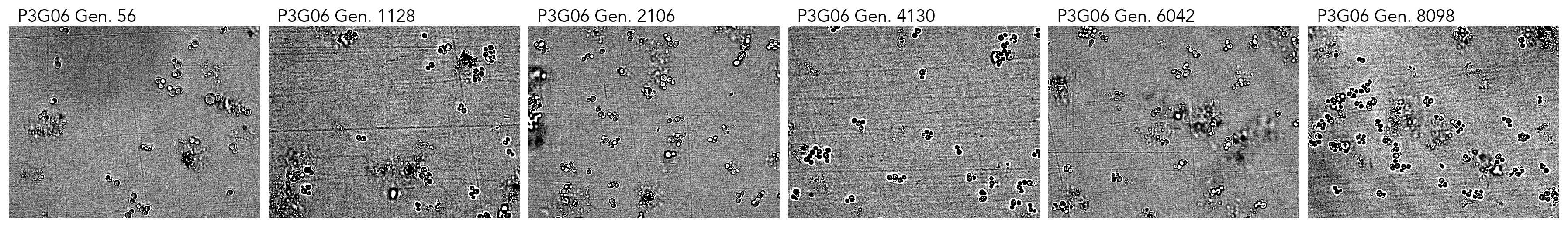

### cropped_P3G09.png

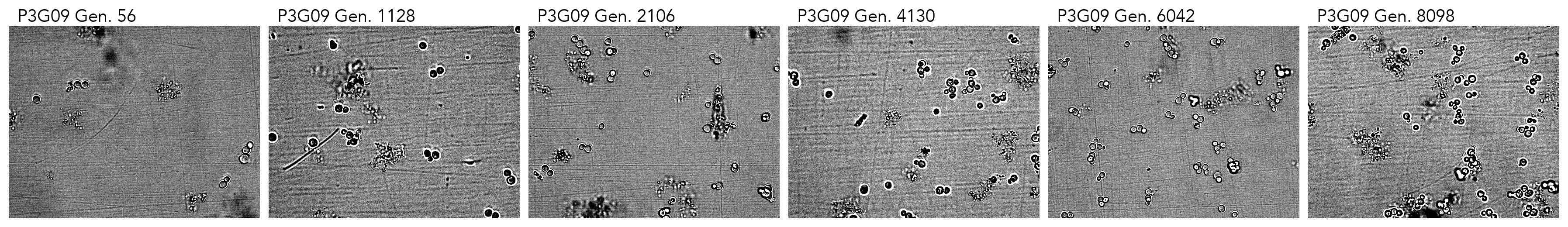

### cropped_P3G10.png

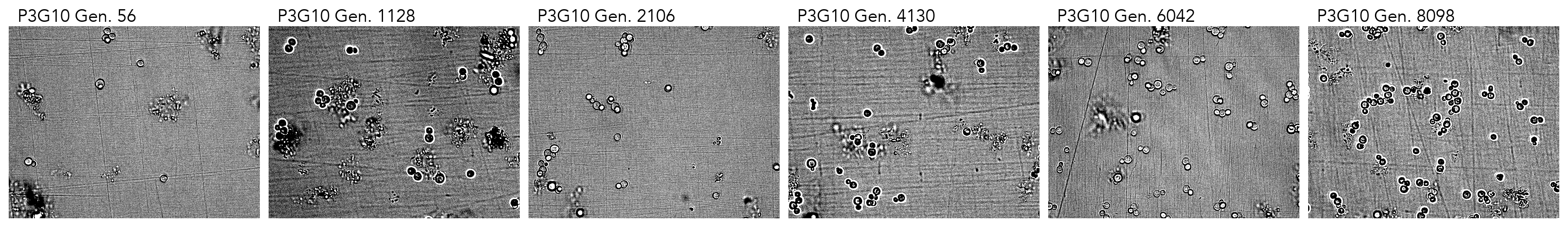

### cropped_P3G11.png

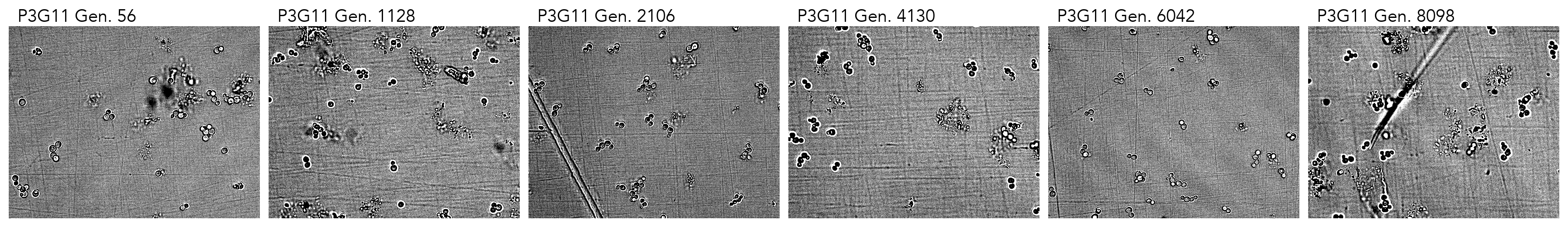
